# The ectomycorrhizal basidiomycete *Laccaria bicolor* releases a GH28 polygalacturonase that plays a key role in symbiosis establishment

**DOI:** 10.1101/2021.09.24.461608

**Authors:** Feng Zhang, Aurore Labourel, Mireille Haon, Minna Kemppainen, Emilie Da Silva Machado, Nicolas Brouilly, Claire Veneault-Fourrey, Annegret Kohler, Marie-Noëlle Rosso, Alejandro Pardo, Bernard Henrissat, Jean-Guy Berrin, Francis Martin

**Affiliations:** State Key Laboratory of Grassland Agro-Ecosystems, Institute of Innovation Ecology, Lanzhou University, Lanzhou, 73000, China; Université de Lorraine, INRA, UMR ‘Interactions Arbres/Microorganismes’, Laboratoire d’Excellence ARBRE, INRA-Grand Est, 54280, Champenoux, France; INRA, Aix-Marseille Université, UMR 1163, Biodiversité et Biotechnologie Fongiques, 13009, Marseille, France; Laboratorio de Micología Molecular, Instituto de Microbiología Básica y Aplicada, Departamento de Ciencia y Tecnología, Universidad Nacional de Quilmes and Consejo Nacional de Investigaciones Científicas y Técnicas (CONICET), Bernal, Provincia de Buenos Aires, Argentina; Aix-Marseille Université, CNRS, IBDM, Marseille, France; CNRS, UMR 7257 & Architecture et Fonction des Macromolécules Biologiques, Aix-Marseille Université, Marseille, France; INRA, USC 1408 AFMB, 13288 Marseille, France; Department of Biological Sciences, King Abdulaziz University, Jeddah, Saudi Arabia; Beijing Advanced Innovation Center for Tree Breeding by Molecular Design, Beijing Forestry University, China

**Author notes:** **Authors for correspondence:** Francis Martin and Feng Zhang.

**Keywords:** apoplastic effector, carbohydrate-active enzymes, cell wall-modifying enzymes, pectins, polygalacturonase

## Abstract

- In ectomycorrhiza, root penetration and colonization of the intercellular space by symbiotic hyphae is thought to rely on the mechanical force that results from hyphal tip growth, enhanced by the activity of secreted cell-wall-degrading enzymes.
- Here, we characterize the biochemical properties of the symbiosis-induced polygalacturonase LbGH28A from the ectomycorrhizal fungus *Laccaria bicolor*. The transcriptional regulation of *LbGH28A* was measured by qPCR. The biological relevance of *LbGH28A* was confirmed by generating RNAi-silenced *LbGH28A* mutants. We localized the LbGH28A protein by immunofluorescence confocal and immunogold cytochemical microscopy in poplar ectomycorrhizal roots.
- qPCR confirmed the induced expression of *LbGH28A* during ectomycorrhiza formation. *L. bicolor* RNAi mutants have a lower ability to establish ectomycorrhiza confirming the key role of this enzyme in symbiosis. The purified recombinant LbGH28A has its highest activity towards pectin and polygalacturonic acid. *In situ* localization of LbGH28A indicates that this endopolygalacturonase is located in both fungal and plant cell walls at the symbiotic hyphal front.
- The present findings suggest that the symbiosis-induced pectinase LbGH28A is involved in the Hartig net formation and is an important determinant for successful symbiotic colonization.

## Introduction

In boreal, mountainous and temperate forest ecosystems, trees rely on ectomycorrhizal associations to obtain nitrogen, phosphorus and micronutrients available in soils (Read *et al*., 2004; Smith & Read, 2008; van der Heijden *et al*., 2015). Ectomycorrhizal fungi establish a symbiotic mutualistic interaction with host rootlets leading to morphogenetic and metabolic changes in both fungal and plant symbionts (Brundrett, 2002; Peterson & Massicotte, 2004). During root colonization, symbiotic hyphae first differentiate to form a pseudoparenchymatous mantle, a structure that encloses root apices (Massicotte *et al*., 1989). Subsequently, the hyphae grow between rhizodermal cells forming an intraradical, apoplastic hyphal network, so-called the Hartig net (Balestrini & Kottke, 2016). The development of the Hartig net is initiated from the innermost layer of the mantle sheath and radial ingress of finger-like hyphae takes place in a broad lobed hyphal front (Balestrini & Kottke, 2016). This labyrinthine hyphal structure is a key feature of the functional ectomycorrhiza. The fusion of fungal and plant cell wall polysaccharides and proteins in the Hartig net produces a common apoplast named the symbiotic interface. This is where the bi-directional translocation of metabolites takes place between the symbionts (Smith & Read, 2008). Differentiation of the Hartig net is linked to subtle biochemical changes of both hyphae and cortical cell surfaces (Balestrini & Kottke, 2016). Localized loosening and swelling, and redistribution of un-esterified pectins, have been reported for plant cell walls at the symbiotic interface (Balestrini *et al*., 1996; Balestrini & Bonfante, 2014; Sillo et al. 2016). In addition, immunocytochemical microscopy has revealed changes in the spatial distribution of fungal cell wall proteins, such as hydrophobins and symbiosis-regulated acidic polypeptides, in the ectomycorrhizal *Pisolithus microcarpus* interacting with *Eucalyptus globulus* (Laurent *et al*., 1999; Tagu *et al*., 2001).

Colonization of the root middle lamella by ectomycorrhizal hyphae is thought to mainly rely on the mechanical force generated by the hydrostatic pressure at the tip of growing hyphae (Peterson & Massicotte, 2004). It has also been proposed that auxins, released by colonizing hyphae, could promote root cell wall loosening (Gay *et al*., 1994ab), thus facilitating hyphal ingress. We have suggested that fungal plant cell wall-degrading enzymes (PCWDEs) could also be involved in the middle lamella colonisation by symbiotic hyphae in *L. bicolor-Populus* ectomycorrhizal roots (Martin *et al*., 2008; Veneault-Fourrey *et al*., 2014). Similarly, Sillo *et al*. (2016) have shown by using the comprehensive microarray polymer profiling (CoMPP) technology that a localized degradation of pectin occurs during root colonization in *Tuber melanosporum–Coryllus avellana* ectomycorrhizas. We recently reported that the symbiosis-induced ß-1,4-endoglucanase LbGH5-CBM1 of the ectomycorrhizal fungus *Laccaria bicolor* is a secreted fungal endocellulase acting on poplar cell walls and this enzyme is an important determinant for successful symbiotic fungal colonization (Zhang *et al*., 2018). *LbGH5-CBM1* expression is substantially induced in ectomycorrhizal poplar roots and *L. bicolor* RNAi-silenced mutants with decreased *LbGH5-CBM1* expression have a lower ectomycorrhiza formation rate. *In situ* localization of LbGH5-CBM1 in ectomycorrhizal rootlets shows that this endoglucanase is located in cell walls of hyphae forming the Hartig net and mantle sheath. In addition to *LbGH5-CBM1*, three *L. bicolor* genes coding for pectinases (polygalacturonases) of the glycoside hydrolase family 28 (GH28) are induced in *L. bicolor-Populus* ectomycorrhizal roots (Veneault-Fourrey *et al*., 2014), suggesting that the Hartig net development may require pectin degradation as shown in *T. melanosporum–C. avellana* ectomycorrhizas (Sillo *et al*., 2016).

Pectins are plant polysaccharides, mainly composed of D-galacturonic acid (GalA), present in the middle lamella and in the primary and secondary cell walls, where they accumulate during the early stages of cell expansion. Pectins are classified, based on their chemical composition, in four primary types: homogalacturonan, rhamnogalacturonan I, xylogalacturonan and rhamnogalacturonan II. This linear polymer is formed of 100-200 GalA units linked by α-(1 → 4) glycosidic bonds (Christiaens *et al*. 2016). Polygalacturonases are hydrolytic enzymes that specifically cleave the α (1-4) bonds between adjacent galacturonic acids in pectin polygalacturonates.

In this study, we functionally characterized the ectomycorrhiza-induced endopolygalacturonase LbGH28A of *L. bicolor*. qRT-PCR assay showed that the expression of *LbGH28A* gene is barely detectable in free-living mycelium, while it is strikingly upregulated in ectomycorrhizal root tips. The expression of *LbGH28A* is induced by pectin and polygalacturonic acid suggesting a role in pectin metabolism. Biochemical characterization of the recombinant protein confirmed that LbGH28A is a pectinase. *LbGH28A* RNAi-mutants show reduced mycorrhiza and Hartig net formation, indicating an important role for this gene in the establishment of the symbiotic interface. By indirect immunofluorescence confocal microscopy, we also showed that LbGH28A accumulates at the periphery of hyphae in the Hartig net and mantle. Immunogold cytolocalization by transmission electron microscopy showed that LbGH28A is accumulating in the cell walls at the symbiotic interface. Our findings suggest that the symbiosis-induced LbGH28A pectinase plays a key role in the establishment of ectomycorrhizal symbiosis in concert with other symbiosis-induced PCWDEs.

## Material and Methods

### Biological material and growth conditions

*Laccaria bicolor* (Maire) P.D. Orton, dikaryotic strain S238N, was grown on a low-glucose Pachlewski agar medium (P20) (Pachlewski, 1967), on cellophane membranes placed on the medium surface, at 20°C in the dark. The axenic cultures were sub-cultured for mycorrhiza establishment weekly (Felten *et al*., 2009). Briefly, P20 is a modification of Pachlewski’s medium composed of 0.25 g L^-1^ di-ammonium tartrate, 0.5 g L^-1^ KH_2_PO_4_, 0.25 g L^-1^ MgSO_4_, 1.0 g L^-1^ glucose, 1 mL L^-1^ (diluted 1/10) Kanieltra microelement solution and 20 g L^−^ agar ^1^. To assess the effect of galacturonic acid and pectin on *LbGH28A* expression, *L. bicolor* S238N mycelium was cultured in 100 mL modified Melin Norkrans (MMN) medium for three days before being transferred to a fresh MMN medium containing either 0.5% w/v glucose, polygalacturonic acid (P3889, Sigma-Aldrich, France) or pectin from *Citrus* peel (poly-D-galacturonic acid methyl ester (galacturonic acid ≥74.0 % dried basis)), P9135, Sigma-Aldrich), as the sole carbon source. Cultures were maintained in 250 mL flasks at 20°C in the dark on a rotary shaker (200 rpm) for one week. Then, the mycelium was harvested by filtration through a Büchner funnel under vacuum, washed twice with Milli-Q water, snap frozen in liquid N_2_ and stored at −80°C for further RNA extraction (Dietz *et al*., 2011).

Hybrid grey poplar (*Populus x canescens*, synonym: *Populus tremula* x *alba*, clone INRA 717-1-B4) was used for *in vitro* ectomycorrhizal inoculation. Two-cm-long cuttings were rooted on solid Murashige and Skoog (MS) medium (Felten et al., 2009), for three weeks, and mycelium of *L. bicolor* S238N (wild-type or RNAi-silenced lines) were grown on cellophane membranes on low-glucose Pachlewski agar medium in the dark for 10 days. Membranes with *L. bicolor* mycelium and poplar seedlings with one or two main roots were transferred onto the surface of low-glucose Pachlewski agar medium (0.1% glucose) containing 0.1% MES (M8250, Sigma, France), and covered by another cellophane membrane (i.e., “sandwich technique”) (Felten *et al*., 2009). Inoculated and control plantlets were grown in a controlled environment growth chamber for 16 h (22°C/18°C, day/night) at 50-60% relative humidity and 400 μmol m^−2^ s^−1^ photosynthetic photon flux density. Ectomycorrhizal rootlets, non-mycorrhizal roots and extramatrical hyphæ were sampled three weeks after contact, snap frozen in liquid N_2_ and stored at −80°C until further analyses. The rate of ectomycorrhiza formation was assessed as described in Felten *et al*. (2009).

### Analysis of the *LbGH28A* gene

The gene coding for the predicted polygalacturonase (pectinase) *LbGH28A* of the glycoside hydrolase (GH) 28 family was identified by automatic and manual annotations in the genome of *L. bicolor* S238N-H82 (http://genome.jgi.doe.gov/Lacbi2/Lacbi2.home.html; Joint Genome Institute ID #313960 in v1 and JGI ID #613299 in v2, GenBank accession # XP_001877488.1). The coding sequence of *LbGH28A* was corrected based on Sanger sequencing of its cDNA. Approximately 50 ng of fungal total RNA, extracted using the Qiagen RNAeasy kit (Qiagen, Courtaboeuf, France) from ectomycorrhizal rootlets, was used for the first strand cDNA synthesis following the manufacturer’s instructions (SMARTer RACE cDNA Amplification Kit (Clontech, Cat. No.634923)). The degenerated oligo(dT) and 3’-RACE and 5’-RACE primers used are listed in Supporting Information Table S1. Then, the amplified fragment was cloned in pJET1.2 vector (Thermo Scientific, K1231) and sequenced by Sanger sequencing. The full-length *LbGH28A* cDNA was used as the reference transcript for producing the recombinant protein in yeast.

The nucleotide sequence of the *LbGH28A* gene and its coding sequence was analyzed and compared to sequences deposited in international databases (MycoCosm, GenBank, SwissProt) by using available online tools (https://mycocosm.jgi.doe.gov/mycocosm/home; http://www.ncbi.nlm.nih.gov/, http://www.expasy.org). The putative signal peptide of the LbGH28A protein sequence was predicted using the SignalP 4.1 server (http://www.cbs.dtu.dk/services/SignalP/). The phylogenetic neighbor-joining tree was deduced from the alignment of fungal GH28 proteins using MAFFT (version 7) with the consistency based method and built the tree by using FastTree (version 2.1.11) and MEGAX (version 10.1.8).

### Quantitative RT-PCR

Axenic mycelium of *L. bicolor* S238N (100 mg fresh weight) and 25 to 50 mycorrhizal or non-mycorrhizal lateral rootlets (100 mg fresh weight), sampled on five different root systems of *P. tremula* x *alba* colonized by *L. bicolor*, were ground in liquid nitrogen using a mortar and pestle. Total RNA was extracted using the RNeasy Plant kit (Qiagen) according to the manufacturer’s instructions with the addition of polyethylene glycol 8000 to RLC buffer (25 mg mL ^-1^). To avoid any DNA contamination, a DNA digestion step was performed on-column with DNAse I (Qiagen). RNA quality was checked by using Experion HighSens capillary gels (BioRad, Marnes-la-Coquette, France). Synthesis of cDNA from one μg of total RNA was performed using the iScript kit (#1708891, BioRad, France) according to the manufacturer’s instructions. All primers were ordered from Eurogentec (Angers, France) and PCR amplification was performed using *Taq* DNA Polymerase (Thermo Fisher Scientific, France) according to the manufacturer’s instructions and optimized according to each primer pairing. Real-time PCR analyses were performed using the Fast SYBR Green Master Mix (Applied Biosystems, France) with a final concentration of 0.3 µM of each primer following the manufacturer’s instructions. The thermal-cycling condition parameters of the StepOnePlus System qPCR apparatus (Applied Biosystems, France) were as follows: 95°C for 3 min; 40 cycles of 95°C for 15 s, 60°C for 30 s followed by a melting curve. PCR amplifications were carried out on three biological replicates and included two distinct technical replicates. Transcript abundance was normalized using four constitutively expressed *L. bicolor* genes coding for a histone H4 (JGI ID# 319764), ubiquitin (JGI ID# 446085), a heat shock protein HSP70 (JGI ID# 609242) and a mitochondrial substrate carrier protein (JGI ID# 611151). Primer sequences (Supporting Information Table S1) were designed using the nucleotide sequences retrieved from the *L. bicolor* genome v2.0 (http://genome.jgi.doe.gov/Lacbi2/Lacbi2.home.html) and available online tools (https://www.ncbi.nlm.nih.gov/tools/primer-blast/). Transcript abundance was quantified using the standard curve method of quantification (based on ΔΔCt calculations), as previously described by Pfaffl (2001).

### Generation of transgenic *L. bicolor* strains

*LbGH28A* RNAi was performed using the RNAi/*Agrobacterium*-mediated transformation vector for intron hairpin RNA (ihpRNA) expression, and transformation of *L. bicolor* vegetative mycelium used the pHg/pSILBAγ vector system as described in Kemppainen and Pardo (2010) using cDNA fragments of *LbGH28A*. A 495 bp-sequence (positions 102 to 596) from the coding sequence was targeted for hairpin formation in avoiding the homologous domain shared by other *LbGH28* genes (Supplemental Information Fig. S2). Four randomly selected strains of each *LbGH28A* RNAi-silenced and mock (pHg/empty pSILBAγ vector) transformants were used for further phenotyping. For each image of ectomycorrhizal sections (n = 10), four types of measurements were obtained using FIJI software (https://fiji.sc), including mantle width, root diameter, Hartig net boundary, and root circumference. These measurements were used to calculate both the ratio of average mantle width to average root diameter and average Hartig net boundary to average root circumference.

### Recombinant enzyme production and purification

The methylotrophic yeast *Pichia pastoris* relying on alcohol oxidase (AOX) to metabolize methanol as its sole carbon source was used as a production host. *P. pastoris* expression vector pPICZαA contains the AOX1 promoter for producing heterologous proteins (Ellis *et al*., 1985, Tschopp *et al*., 1987, Koutz *et al*., 1989) and a (His)_6_ tag located at the C-terminus for purification. *P. pastoris* strain X33 was purchased from Invitrogen (Cergy-Pontoise, France). The nucleotide sequence of *LbGH28A* without its signal peptide sequence (Supplemental Information Figure S1) was codon optimized for expression in *P. pastoris* and synthesized by Genscript (NJ, USA). The synthesized sequence was then inserted into the expression vector pPICZαA in frame with the yeast α-factor secretion peptide at the N-terminus and the (His)_6_ tag at the C-terminus, and under the control of the *AOX1* promoter. The expression protocol is described in Couturier *et al*. (2011). Large scale production of the protein was performed in 2 L non-baffled flasks, each containing 500 mL of BMGY medium (1% yeast extract, 2% peptone, 1% glycerol, 400 μg l^-1^ biotin, and 0.1 M potassium phosphate, pH 6.0). *P. pastoris* was grown overnight at 30°C at 200 rpm, and recovered by centrifugation the following day when the absorbance was between two and six units. Pellets from five flasks were pooled and resuspended in 100 mL of BMMY medium (1% yeast extract, 2% peptone, 400 μg l^-1^ biotin, 1% methanol, and 0.1 M potassium phosphate, pH 6.0) for each 500-mL flask. Induction was carried out for three days with the addition of three ml methanol per flask per day. The supernatant was then collected and after setting the pH to 7.8 with NaOH 1M, it was filtered through a 0.22 μm filter membrane (Durapore GV membrane filters, 0.22 μm, Millipore, Molsheim, France). A HisTrap HP column (5 ml, GE Healthcare, France), prepacked with Ni High Performance Sepharose, was connected to an ÄKTA purifier chromatography system (GE Healthcare, France) and equilibrated with the equilibration buffer (50 mM Tris-HCl pH 7.8, 150 mM NaCl, 10 mM imidazole) before purification according to the instructions of the manufacturer. The protein was eluted with 50 mM Tris-HCl, pH 7.8, 150 mM NaCl and 250 mM imidazole. Elution was monitored by measuring the absorbance at 280 nm. The fractions corresponding to the eluted protein were pooled, and loaded onto an ultrafiltration column (Vivaspin 3 or 10 kDa MWCO, PES, Sartorius, Palaiseau, France) for concentration and buffer exchange at 4 °C. The proteins were stored at 4 °C in a 50 mM sodium acetate buffer, pH 5.2. The concentration of the pure proteins was determined by measuring the absorbance of the solution at 280 nm on a Nanodrop 2000 (Thermo Fisher Scientific, France) and calculated using Beer’s law and the extinction coefficient of the protein as determined by ProtParam (http://web.expasy.org/protparam/).

### Enzymatic activity assay

The enzyme activity of LbGH28A was assayed in triplicate by measuring the release of reducing sugars ends from pectin and quantified using the 3-amino-5-nitrosalicylic acid (DNS) method (Couturier *et al*., 2011). LbGH28A was incubated with the following polysaccharides: pectin of citrus peel (P9135, Sigma Aldrich, France), polygalacturonic acid (Megazyme, Wicklow, Ireland), rhamnogalacturonan (R3875, Sigma Aldrich, France), potato pectic galactan (Megazyme) and lupin galactan (Megazyme). Briefly, 1.53 µg of enzyme was mixed with 0.5% (w/v) substrate in 50 mM sodium acetate buffer at pH 4 and incubated at 40°C for 10 min in a total volume of 110 µl. Then, the release of reducing sugars ends was determined using the 3,5-dinitrosalicylic acid (DNS) assay (Sigma). Sixty μl of DNS were added to 60 µl of the enzymatic reaction and incubated at 100 °C for 10 min, and the absorbance at 540 nm was measured. One unit of enzymatic activity (U) is defined as the amount of enzyme required to release one µmol of glucose reducing-sugar equivalents per minute (µmol.min^-1^) under the defined assay conditions.

For optimal pH and temperature determination, polygalacturonic acid, 0.5% (w/v), was used as substrate. The pH optimum was determined as described above using 50 mM sodium acetate with different pH (3.5, 4.0, 4.5, 5.0 and 5.5) for 10 min at 40°C. For optimal temperature determination, enzyme activity was measured at 30°C, 35°C, 40°C, 45°C, 50°C and 55°C for 10 min.

### Confocal microscopy and indirect immunofluorescent localization

Three-week-old ectomycorrhizal rootlets from grey poplar or free-living *L. bicolor* S238N mycelium were fixed for eight hours in 4% (w/v) paraformaldehyde in phosphate-buffered saline (PBS) buffer (137 mM NaCl, 2.7 mM KCl, 10 mM Na_2_HPO_4_, 1.8 mM K_2_HPO_4_, pH 7.4) at 4°C. The root segments were embedded in agarose 5% (w/v) and cut into 25-30 μm radial sections with a VT1200S vibratome (Leica Microsystems, Nanterre, France). Radial sections were sampled from 15 ectomycorrhizal rootlets (at 200 µm from the root apex) to assess LbGH28A protein accumulation. Sections were retrieved with a brush and carefully transferred onto watch glasses and then were stained according to Felten *et al*. (2009).

A solution containing 3.5 mg of purified recombinant LbGH28A protein was used to elicit the production of polyclonal antibodies in rabbit according to the manufacturer’s procedure (Eurogentec, Seraing, Belgium). The indirect immunofluorescent (IIF) localization of the LbGH28A protein was performed by confocal microscopy as described in Zhang *et al*. (2018). Briefly, sections were transferred onto watch glasses and incubated in 1% BSA for one hour. Then, BSA was removed and sections were incubated overnight with purified anti-LbGH28A protein rabbit antibody diluted 1:1,000 in PBS containing 0.5% (w/v) BSA at 4°C. The segments were then washed five times in PBS and incubated in the secondary antibody conjugate, a 1:500 dilution of goat anti-rabbit IgG-AlexaFluor 488 conjugate (A-11008, ThermoFisher, France) in PBS for 2 h. After five more washes in PBS, sections were mounted in 80% glycerol (Merck), 20% PBS, 5% w/v propyl gallate (Fluka) and viewed by a Zeiss LSM 800 microscope equipped with X10, X40, numerical aperture 1.4. The excitation and emission wavelengths for the Alexa Fluor 488 dye were 500 to 550 nm, respectively. Optical sections were collected at 0.1 to 0.7 mm intervals with Kalman averaging. As a control, sections were incubated with IgG purified from pre-immune serum diluted to the same concentration as anti-LbGH28A IgG. In addition, we performed a competition epitope binding assay by incubating sections with both anti-LbGH28A protein rabbit antibody diluted 1:1,000 and increasing concentration of recombinant LbGH5-CBM1 protein in PBS containing 0.5% (w/v) BSA at 4°C.

### Ultrastructural localization by immunogold labelling

Three-week-old ectomycorrhizal root tips or axenic mycelium of *L. bicolor* S238N were sampled and fixed with 2.5 % glutaraldehyde, 2 % paraformaldehyde in saline phosphate buffer (PBS) for 2 h. The samples were washed in PBS and post-fixed in 1 % osmium tetroxide in distilled water for 1 h, washed and incubated in uranyl acetate 1 % in distilled water overnight at 4°C. Samples were then dehydrated in 100 % ethanol and acetone, and embedded in Epon resin. Ultrathin sections (90 nm) were performed on a UC7 Leica Ultra microtome (Leica, Netherlands). Sections on nickel grids were incubated with saturated sodium metaperiodate for 3 min. The grids were washed for 4 min in TBS with 1 %Triton X-100 and then incubated in 10 % normal serum in TBS for 30 min, followed by an overnight incubation with rabbit anti-LbGH28A antibodies (dilution 1/5) at 4°C. The grids were washed in saline Tris buffer (TBS), incubated with secondary antibodies (6 nm-gold particles linked to anti-rabbit antibodies, Aurion, dilution 1/15) for one hour at 37°C, then washed again in TBS. Grids were incubated for 10 min in 2.5 % glutaraldehyde in 0.05 M cacodylate buffer, washed in water and counter-stained with 1 % uranyl acetate (for 5 min) and lead citrate (for 2 min), respectively. Acquisitions were performed on a Tecnai G2 electron microscope (FEI, Netherlands) at 200 kV. Micrographs were acquired with a Veleta camera (Olympus, Japan) on triplicate samples.

## Results

### The symbiosis-induced polygalacturonase LbGH28A from L. bicolor

Among the PCWDE genes previously reported as induced in *L. bicolor–P. tremula x alba* ectomycorrhizal rootlets (Veneault-Fourrey *et al*., 2014), three genes are annotated as pectinases from the GH28 family. These GH28 pectinases are predicted to hydrolyze polygalacturonates, the major component of plant pectins. Gene # 613299 belongs to the endo-polygalacturonase GH28 subfamily A (Kohler *et al*., 2015) and has thus been named *LbGH28A*. It is induced during the earlier steps of *L. bicolor–P. tremula x alba* symbiosis suggesting it may play a role in plant cell wall remodeling during the Hartig net formation (Veneault-Fourrey *et al*., 2014). However, the enzymatic activity of LbGH28A has not been confirmed so far. Here, we functionally characterized this *LbGH28A* gene. The gene contains 2,017 nt with 18 introns, the predicted full-length transcript is 1,065 nt and the deduced polypeptide sequence contains 361 amino-acids (molecular mass (MM) of 37.2 kDa, pI 7.61). The LbGH28A polypeptide contains a predicted signal peptide (positions 1 to 16) and GH28 (endo-polygalacturonase) catalytic motif (positions 36 to 344, Pfam 00295) (Supplemental Information Fig. 1). To validate the JGI gene annotation, the full-length *LbGH28A* cDNA was amplified by RACE PCR from total RNA extracted from ectomycorrhizal rootlets and sequenced (Supplemental Information Fig. 1). The ortholog from *L. amethystina* (JGI ID #671067, LaGH28A) is highly similar to LbGH28A (89% similarity, Supplemental Information Fig. S2), indicating that this symbiosis-induced polygalacturonase is highly conserved within the *Laccaria* genus. LbGH28A is also orthologous to endopolygalacturonases of the ectomycorrhizal *Cortinarius glaucopus*, the wood decayer *Hypholoma sublateritium* and litter decayer *Agrocybe pediades* (Fig. 1b, see also Supplementary Fig. 10 in Kohler *et al*., 2015). Interestingly, it shows a low protein sequence similarity (49.5 %) to the symbiosis-induced polygalacturonase of the ascomycetous *T. melanosporum*.

**Figure 1.**
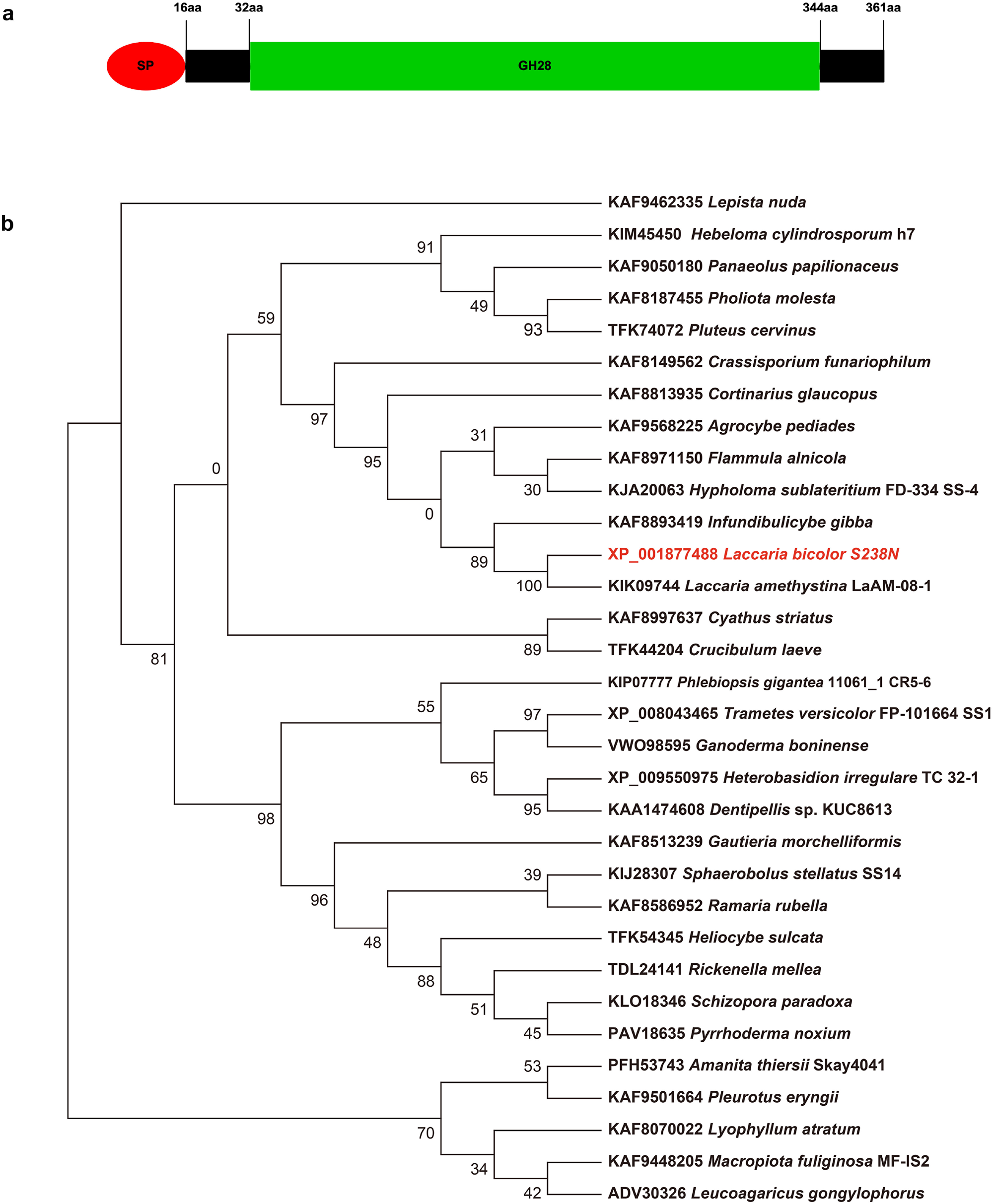
The symbiosis-induced endopolygalacturonase LbGH28A from *L. bicolor*. (**a**) LbGH28A modular structure showing the putative signal peptide (in pink) and the catalytic GH28 domain (in green). (**b**) Phylogenetic relationship of LbGH28A to other related GH28 polygalacturonases in Agaricomycotina. The phylogenetic neighbor-joining tree was deduced from the alignment of fungal GH28 proteins using MAFFT (version 7) with the consistency-based method. The tree was constructed using FastTree (version 2.1.11). and MEGAX (version 10.1.8).

### *LbGH28A* expression is induced by pectin and upregulated in symbiosis

*LbGH28A* is a pectin-inducible gene as the presence of *Citrus* pectin in the growth medium increased its transcription in free-living mycelium by comparison to glucose (Fig. 2A). Polygalacturonic acid also induced the *LbGH28A* expression, but this was not statistically significant. We confirmed by qPCR that *LbGH28A* is transcribed at a lower, constitutive level in the free-living mycelium of *L. bicolor* S238N grown on Pachlewski agar medium (containing 5.5 mM glucose), while it is induced 2.27-fold in ectomycorrhizal hyphae after three weeks post contact (Fig. 2b). *LbGH28A* transcripts were not detected in fruiting bodies of *L. bicolor* (data not shown).

**Figure 2.**
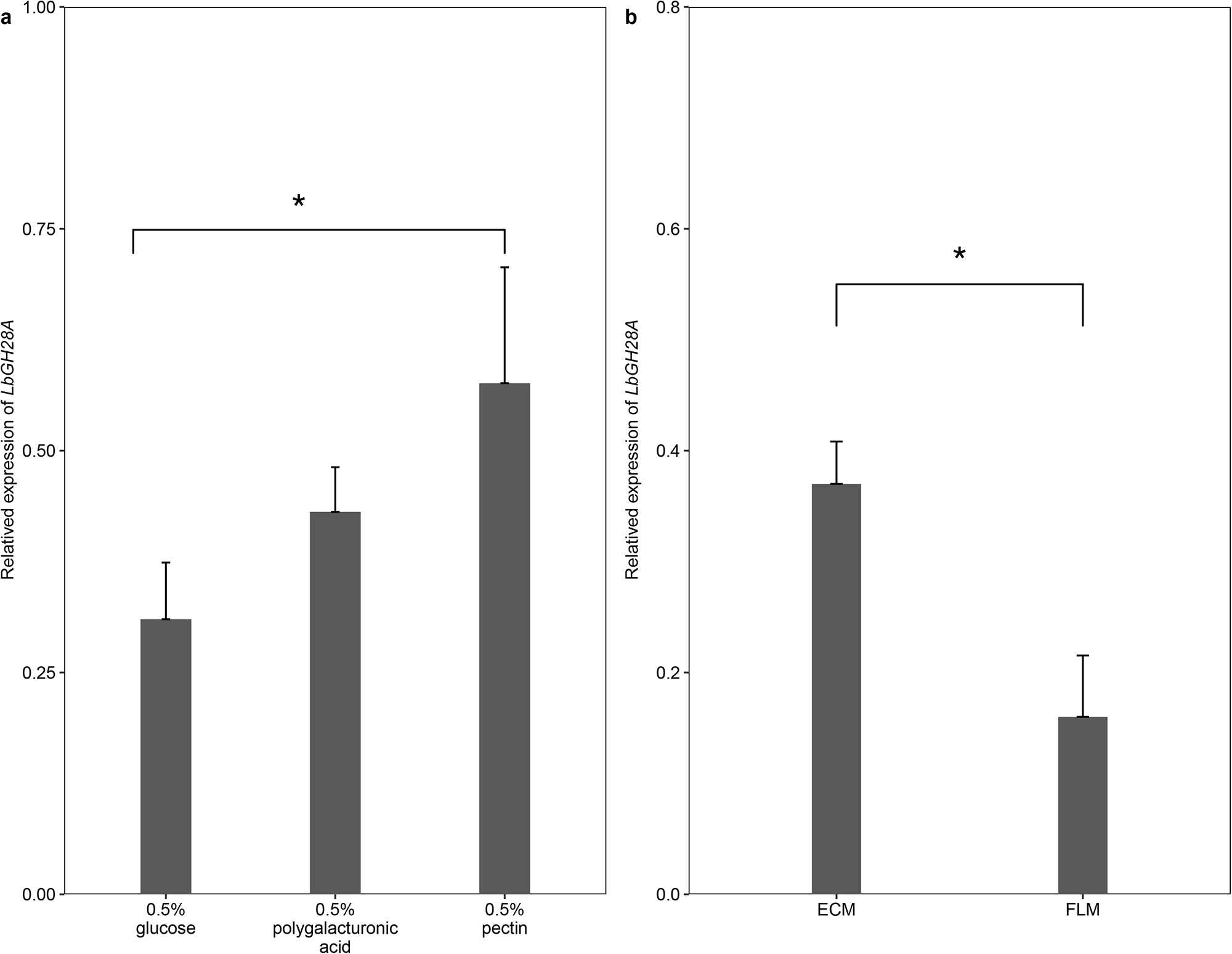
*LbGH28A* expression is induced by polygalacturonate and pectin, and upregulated in ectomycorrhiza. (**a**) *LbGH28A* relative expression in *L. bicolor* free-living mycelium grown on 0.5% glucose, 0.5% polygalacturonic acid or 0.5% *Citrus* pectin as a sole source of carbon. (**b**) Relative *LbGH28A* expression in wild-type S238N (WT) in free-living mycelium and ectomycorrhizal root tips assayed by quantitative RT-PCR. Data presented are the means of three independent biological replicates with two technical replicates per each fungal strain. Error bars represent standard deviation (± SD).

### *LbGH28A* expression is critical for the formation of the Hartig net

To confirm the role of the *LbGH28A* gene in ectomycorrhiza establishment, we generated a series of transgenic *L. bicolor* lines with a lowered level of *LbGH28A* by knocking down its expression by RNAi-mediated gene silencing. Four randomly selected *L. bicolor* mutant strains (*LbGH28* RNAi-1, *LbGH28* RNAi-2, *LbGH28* RNAi-3, *LbGH28* RNAi-4) and empty vector controls (EV7 and EV9) were co-cultured with poplar seedlings to establish ectomycorrhiza. The *LbGH28* RNAi-3 and *LbGH28* RNAi-4 mutants were significantly impaired in their ability to form ectomycorrhizas (Fig. 3a) by comparing to empty vector controls. While the percentage of ectomycorrhizal root tips was 57 % and 53 % for the EV controls, it reached only 11 % and 16 % for the *LbGH28* RNAi-3 and *LbGH28* RNAi-4 mutants, respectively. The rate of ectomycorrhiza formation by the *LbGH28* RNAi-1 (53 %) and *LbGH28* RNAi-2 (50 %) transformant lines was similar to the EV controls. To assess the development of symbiotic structures *in planta*, we harvested colonized root tips and measured the mantle sheath thickness and the Hartig net depth after staining of hyphae by using wheat germ agglutinin (WGA) conjugated with Alexa Fluor 488 (Fig. 4). The laser-scanning confocal microscopy images showed that the *LbGH28A* RNAi mutant lines have a significantly reduced mantle sheath and Hartig net (Fig. 3c,d). The mantle thickness was 19.5 ± 0.9 µm and 18.2 ± 2.6 µm per root section for the empty vector controls, while it ranged from 10.4 ± 0.7 µm to 21.7 ± 1.3 µm for the *LbGH28A* RNAi mutant lines. These values were significantly lower for the *LbGH28* RNAi-2, *LbGH28* RNAi-3 and *LbGH28* RNAi-4 mutants (P<0.01, student’s test, n>150), despite the substantial heterogeneity observed between ectomycorrhizal root tips. The Hartig net depth was 8.9 ± 0.2 µm and 11.6 ± 0.5 µm per root section for the empty vector controls, whereas it was significantly lower for the *LbGH28A* RNAi mutant lines (P<0.01, student’s test, n>150), ranging from 4.4 ± 0.2 µm to 8.9 ± 0.4 µm. Taking together, these morphometric measurements suggest that *LbGH28A* facilitates the fungal ingress into the host intercellular space and as a consequence, participates to the symbiosis establishment.

**Figure 3.**
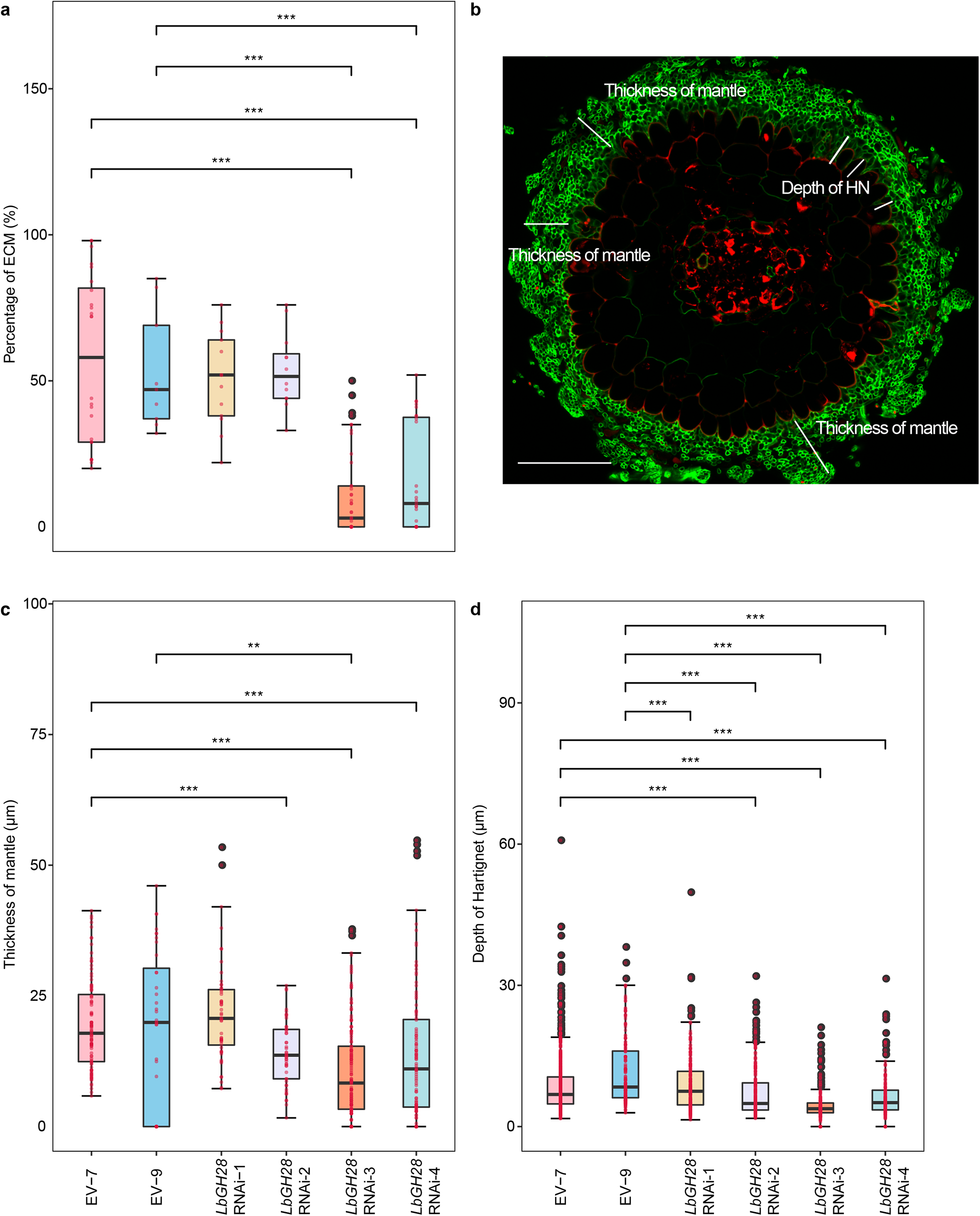
Rate of formation and morphometric features of ectomycorrhizas established by *LbGH28A* RNAi-silenced- and control strains. **(a)** Percentage of ectomycorrhizal rootlets formed by the empty vector control strains (EV-7, EV-9) and RNAi-silenced lines (*LbGH28* RNAi-1, *LbGH28* RNAi-2, *LbGH28* RNAi-3, *LbGH28* RNAi-4) three weeks after contact. For each image of ectomycorrhizal sections (b), morphometric measurements (i.e., Hartig net depth (c) and mantle sheath thickness (d)) were carried out using the FIJI software (https://fiji.sc). (b) A section of ectomycorrhizal roots showing how the mantle sheath thickness and Hartig net depth were measured. Values are shown for the rate of ectomycorrhiza formation (a), the mantle sheath thickness (c) and the Hartig net depth (d). Bars represent the mean of the data and error bars represent the SE of the mean. **, *P* <0.01, ***, *P* <0.001, student’s *t* test.

**Figure 4.**
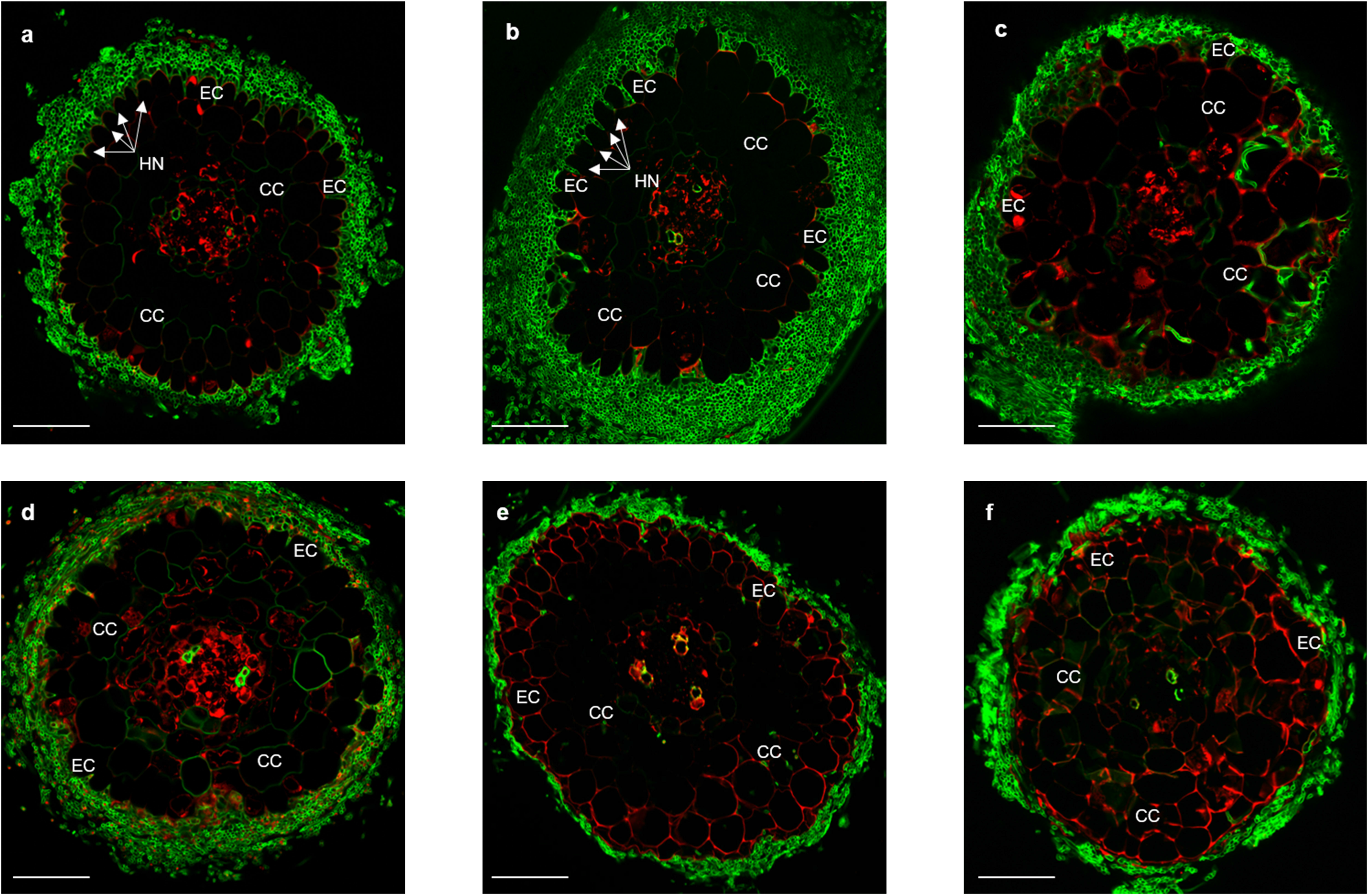
*LbGH28A* is required for Hartig net formation. Representative transverse cross sections of *Populus* roots colonized by (**a-b**) empty vector (EV-7 and EV-9) control strains or (**c-f**) RNAi-silenced (*LbGH28* RNAi-1, *LbGH28* RNAi −2, *LbGH28* RNAi −3, *LbGH28* RNAi −4) strains sampled three weeks after contact. Colonized roots were sectioned and stained with WGA conjugated with Alexa Fluor 488 (green) and propidium iodide (red) and imaged on a confocal laser scanning microscope. M, Mantle; HN, Hartig net; EC, epidermal cells; CC, cortical cells. Scale bar, 50 μm

### Cloning, expression and purification of LbGH28A in *P. pastoris*

LbGH28A, without its native signal peptide, was successfully expressed in *P. pastoris* and purified by affinity chromatography enabling its biochemical characterization. SDS-PAGE showed that the recombinant LbGH28A migrated as a single band with a molecular mass (MM) of 55 kDa (Fig. 5a). This MM was higher than the predicted MM (37.2 kDa) suggesting that the mature protein synthesized in yeast was glycosylated. The purified recombinant LbGH28A protein was then used to assess the enzyme activity and produce polyclonal antibodies in rabbit.

**Figure 5.**
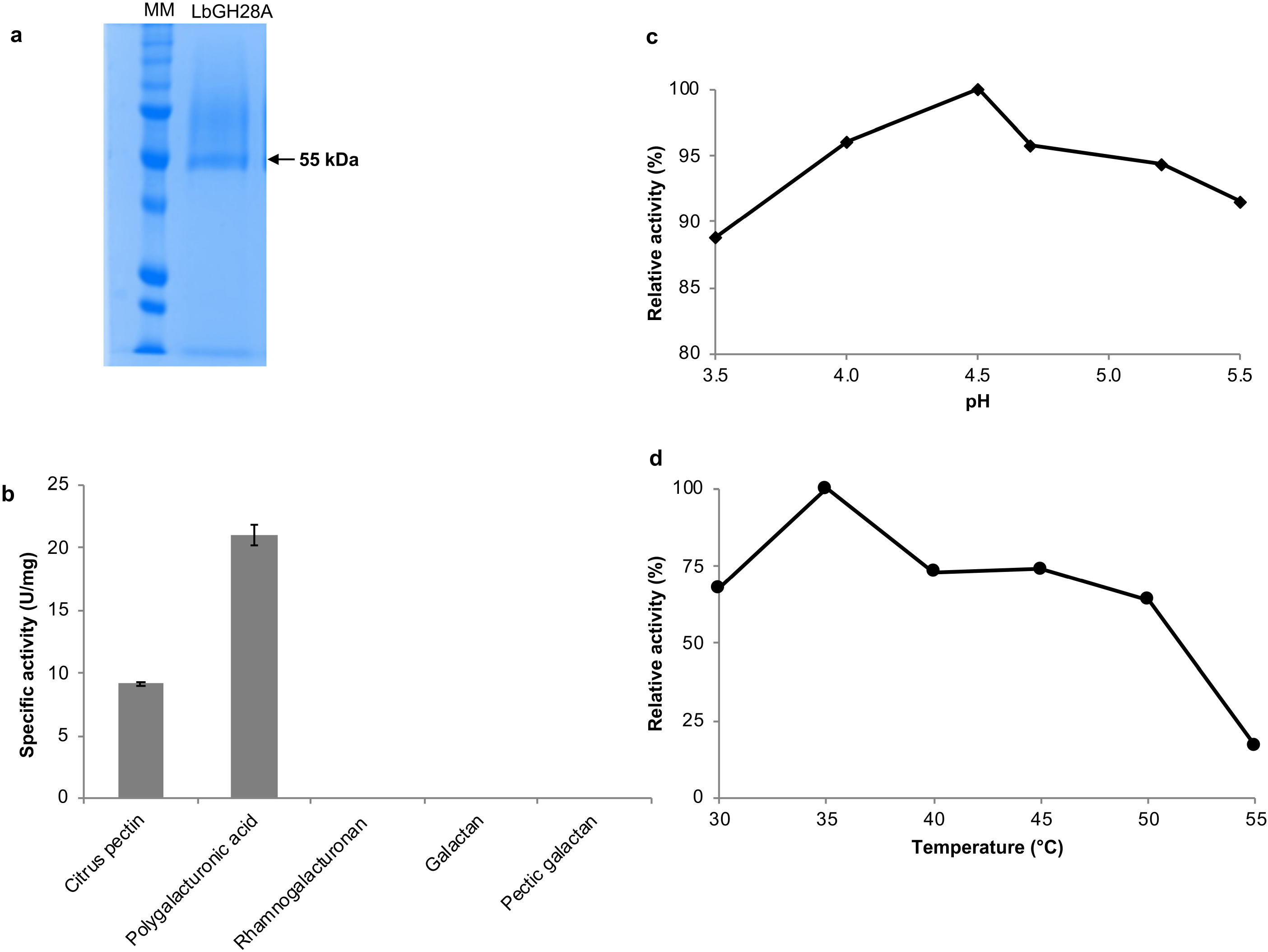
Heterologous production and enzymatic activity of LbGH28A. **(a)** SDS-PAGE analysis of the purified recombinant protein LbGH28A (5 µg). (**b**) Specific enzyme activity on different substrates. (**c**) Effect of pH and (**d**) temperature on LbGH28A specific enzyme activity. Hydrolysis of polygalacturonic acid (5 mg ml^-1^) by the recombinant LbGH28A was measured in 50 mM sodium acetate buffer for 10 min. MM, molecular mass markers (15 to 180 kDa). Data presented are the means of three independent replicates. Error bars represent standard deviation (± SD).

### Enzymatic properties of LbGH28A

The substrate specificity of the recombinant LbGH28A was assayed by using *Citrus* pectin, polygalacturonic acid, rhamnogalacturonan, galactan or pectic galactan. LbGH28A exhibits an endo-polygalacturonase activity, as indicated by the increase of released reduced sugars from *Citrus* pectin and polygalacturonic acid (Fig. 5b). No activity was observed on rhamnogalacturonan, galactan or pectic galactan (Fig. 5b). The optimal pH and temperature of LbGH28A are 4.5 and 35°C, respectively (Fig. 5c,d).

### LbGH28A is localized on fungal and plant cell walls

We performed immunolabelling experiments on *L. bicolor–P. tremula x alba* ectomycorrhizal rootlets to localize LbGH28A. Probing ectomycorrhizal root sections with anti-LbGH28A antibodies led to an intense labeling of hyphae constituting the mantle and the Hartig net (Fig. 6a). Labeling was mainly detected at the periphery of the hyphae and coincided with the cell wall chitin labeled by wheat germ agglutinin (WGA633), supporting a cell wall and/or apoplastic localization. In the presence of the recombinant LbGH28A (competitive assay), the binding of antibodies to LbGH28A was precluded and no signal was detected, confirming the high specificity of the immune serum (Fig. 6b). No labelling was detected when a pre-immune serum was used instead of LbGH28A-antibody (Fig. 6c).

**Figure 6.**
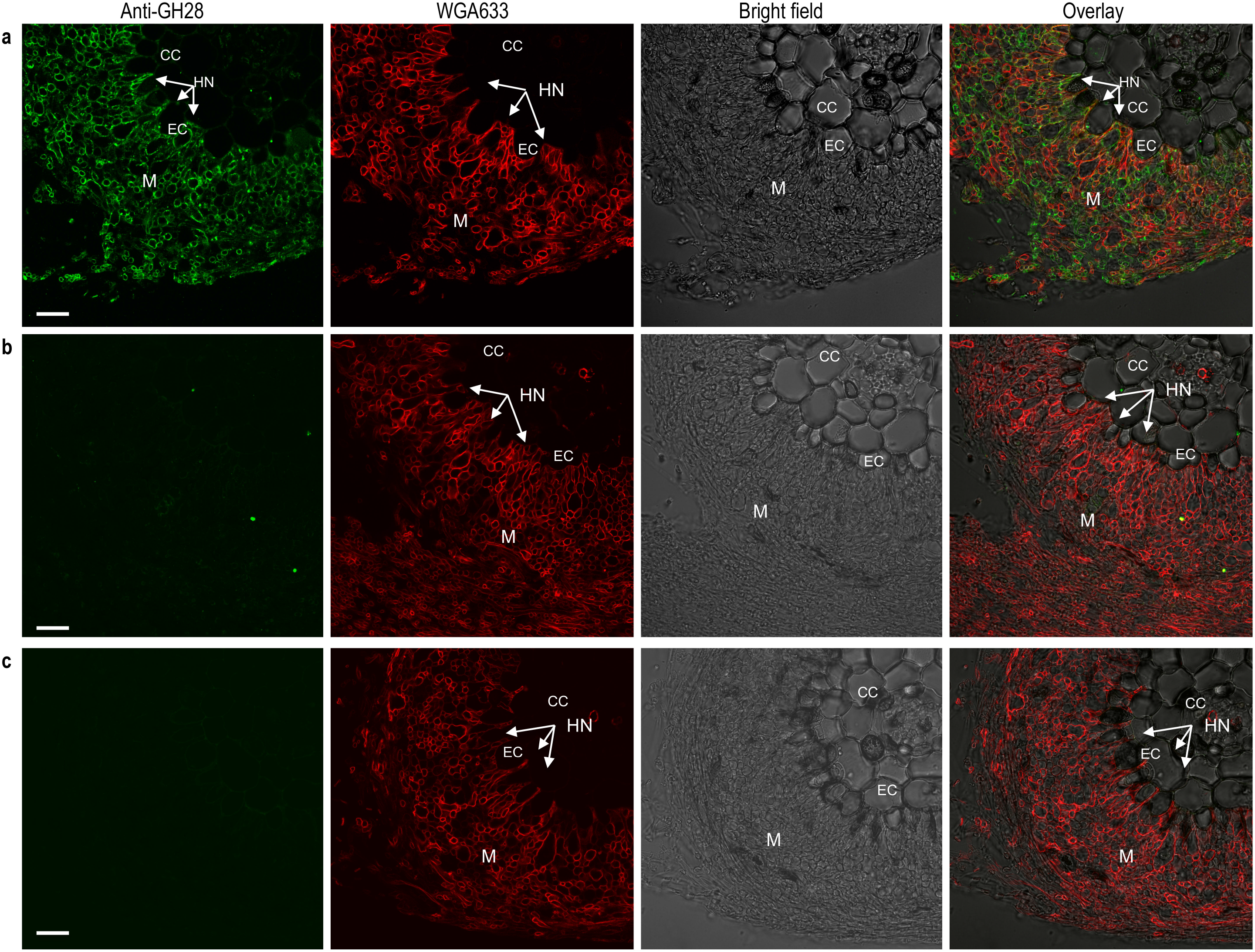
Immunolocalization of LbGH28A in *Populus tremula x alba–L. bicolor* ectomycorrhizal rootlets. All images were obtained by using indirect immunofluorescence confocal laser microscopy. (**a**) Transversal sections of 3-week-old ectomycorrhizas stained for LbGH28A with anti-LbGH28A immune serum (green), for fungal cell wall chitin with WGA633 (red), bright field and the overlaid images. (**b**), A pre-immune rabbit serum was used instead of anti-LbGH28A immune serum. (**c**) Binding of anti-LbGH28A antibodies to LbGH28A was blocked by pre-incubating sections in the presence of the recombinant LbGH28A protein confirming the binding specificity. Abbreviations: M, fungal mantle; HN, Hartig net; EC, epidermal cell; CC, cortical cell. Scale bar, 20 μm.

Immunocytolocalization of LbGH28A was performed on ultrathin sections of *L. bicolor* free-living mycelium and *L. bicolor–P. tremula x alba* ectomycorrhizal rootlets by transmission electronic microscopy (TEM) (Fig. 7). At the tip of the symbiotic hyphae penetrating between epidermal cells, the 6 nm-gold particles, indicating the presence of LbGH28A, were detected in the merged fungal and plant cell walls of the symbiotic interface (Fig. 7cd). This localization confirms that LbGH28A is secreted in the symbiotic interface where it bounds to fungal and plant cell wall materials. No labelling was detected when a pre-immune serum was used instead of LbGH28A-antibody (Fig. 7ef). In addition, no labelling was observed in the plant cell walls of non-mycorrhizal poplar rootlets when pre-immune and immune sera were used (Fig. 7ghij), confirming that the anti-LbGH28A antibody is specific to the fungal LbGH28A with no cross-hybridization to poplar polygalacturonase(s). LbGH28A was also detected in the cell walls of the free living mycelium (Fig. 7ab), despite the low expression of LbGH28A (Fig. 2b). Finally, we also immunolocalized the symbiosis-induced ß-1,4-endoglucanase LbGH5-CBM1 from *L. bicolor* (Zhang *et al*., 2018) in the ectomycorrhizal root tips used for the LbGH28A immunolocalization. LbGH5-CBM1 was strictly located in fungal cell walls (Fig. 8).

**Figure 7.**
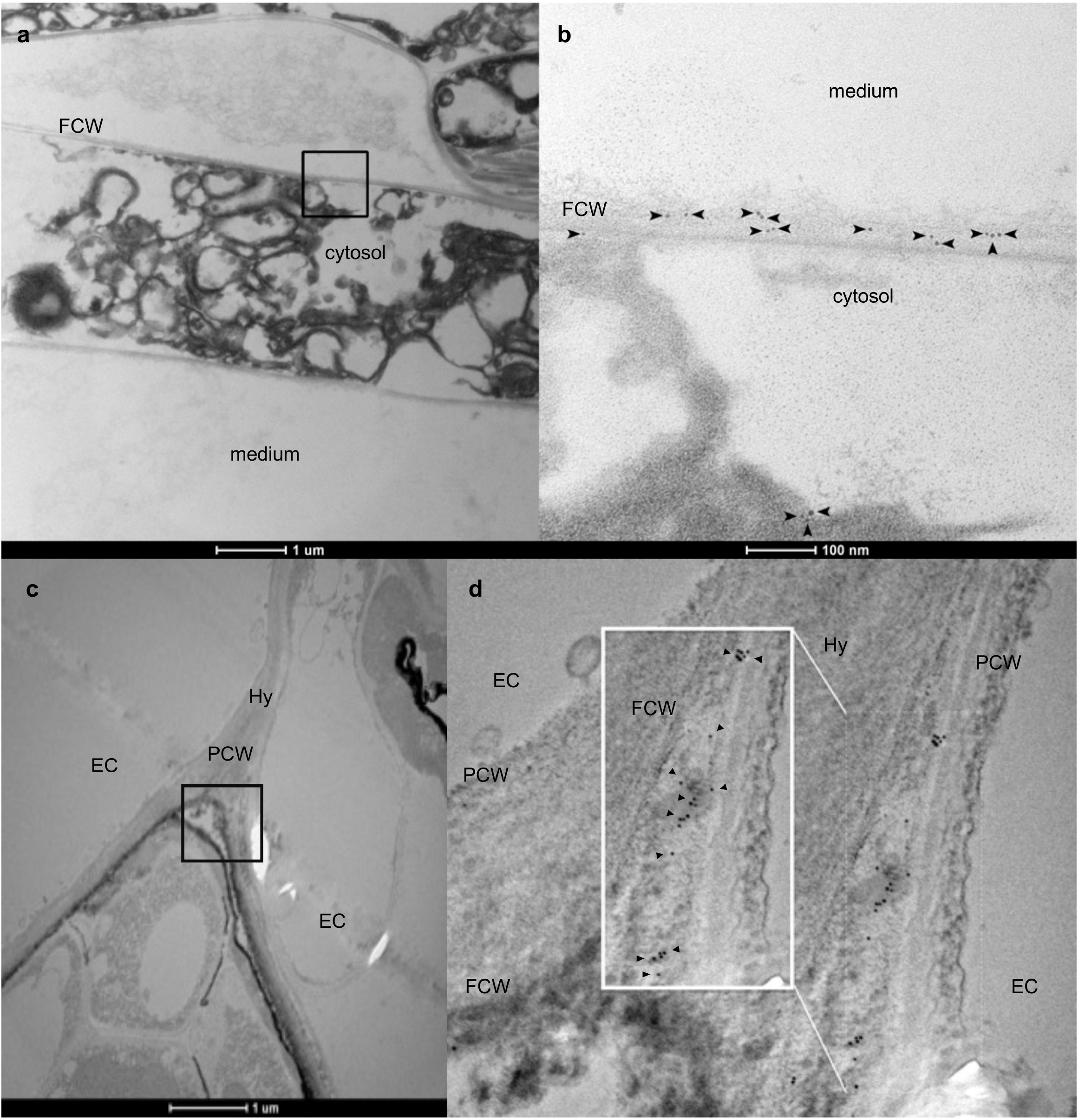
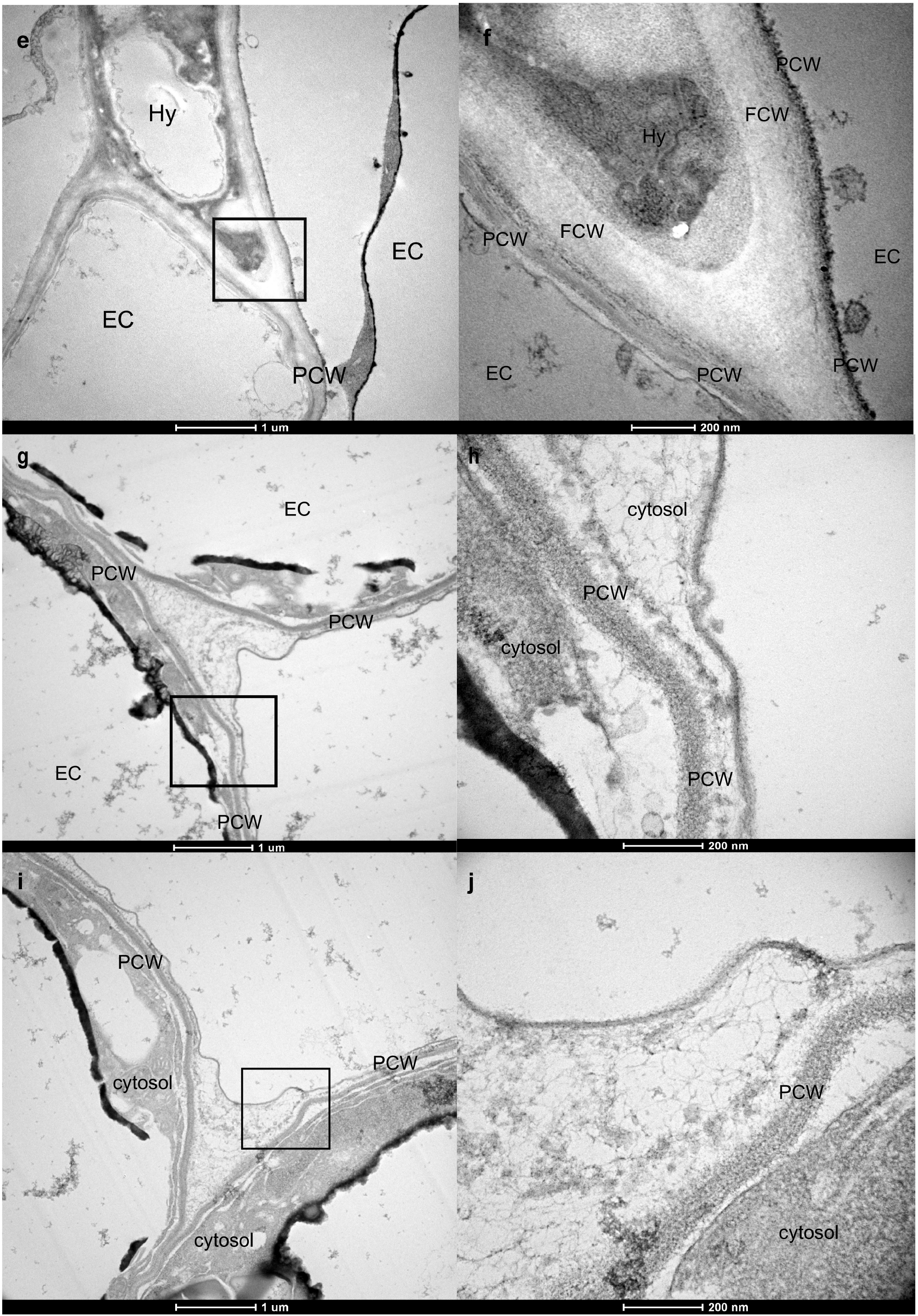
Immunogold cytolocalization of LbGH28A at the symbiotic interface of ectomycorrhizal root tips. TEM images showed the 6 nm-gold particles corresponding to the localization of the LbGH28A antibody/LbGH28A protein in the free-living mycelium of *L. bicolor* (**a, b**) or *L. bicolor* hyphal tips (HT) colonizing the middle lamella/apoplastic space (AS) between two epidermal cells (EC) of *P. tremula x alba* roots (**c, d**). No labelling was observed when sections were incubated without antibody (**e, f**); no labelling was observed in cell walls of non-mycorrhizal rootlets incubated with LbGH28A antibody (**g, h**) or without antibody as control (**i, j**). Ultra-thin transverse sections of 3-week-old ectomycorrhizas or poplar roots were incubated with anti-LbGH28A rabbit antibodies coupled to 6 nm gold particles. Gold particles indicating LbGH28A localization are highlighted by black arrows. PCW; plant cell wall, FCW; fungal cell wall, Hy, fungal hyphae.

**Figure 8.**
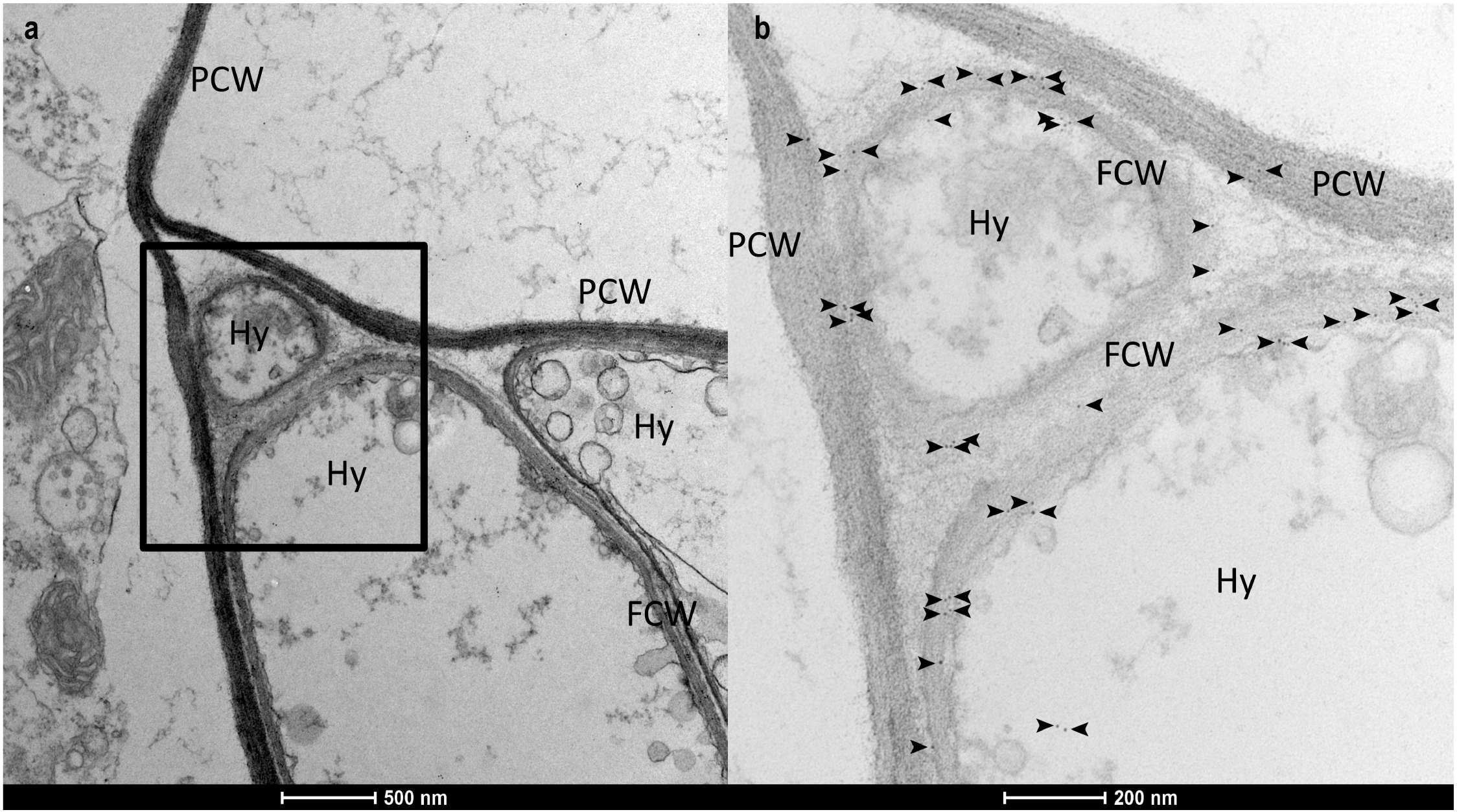
Immunogold cytolocalization of LbGH5-CBM1 in ectomycorrhiza and free-living mycelium of *L. bicolor*. TEM images showing the localization of the LbGH5-CBM1 antibody/LbGH5-CBM1 protein in *L. bicolor* hyphal tips (Hy) colonizing the middle lamella/apoplastic space between two epidermal cells of *P. tremula x alba* roots. Ultra-thin transverse sections of 3-week-old ectomycorrhizas were incubated with anti-LbGH5-CBM1 rabbit antibodies (Zhang *et al*., 2018) coupled to gold particles. Gold particles indicating LbGH5-CBM1 localization are highlighted by black arrows. PCW; plant cell wall, FCW; fungal cell wall, Hy, fungal hyphae.

## Discussion

Colonization of the root middle lamella space by ectomycorrhizal hyphae is required for the formation of the Hartig net, the symbiotic interface, and for subsequent symbiosis development. This fungal penetration between epidermal and cortical root cells leads to biochemical changes in both hyphal and root cell walls (Balestrini & Kottke, 2016). These changes include localized loosening and swelling of cell walls, and redistribution of un-esterified pectins (Balestrini *et al*., 1996; Balestrini & Bonfante, 2014; Sillo et al., 2016). Fungal ingress in the middle lamella is thought to rely on the mechanical force that results from hyphal tip growth (Peterson & Massicotte, 2004), the effect of auxins released by colonizing hyphae (Gay *et al*., 1994 a,b) and the activity of fungal PCWDEs (Veneault-Fourrey *et al*., 2014; Sillo *et al*., 2016). Although the arsenal of PCWDEs encoded by *L. bicolor* and other ectomycorrhizal fungi is restricted by comparison to saprotrophic fungi (Martin *et al*., 2008; Kohler *et al*., 2015; Miyauchi *et al*., 2020), several of the remaining PCWDE genes are induced in ectomycorrhizal associations (Martin *et al*., 2008, 2010; Kohler *et al*., 2015; Peter *et al*., 2016; Sillo *et al*., 2016; Miyauchi *et al*., 2020). Among the >20 symbiosis-upregulated PCWDEs of *L. bicolor*, endoglucanases (GH5), pectin methyl esterases (CE8), polygalacturonases (GH28) and cellulose acting-lytic polysaccharide monooxygenases (LPMO, AA9) are induced at different stages of ectomycorrhiza establishment (Veneault-Fourrey *et al*., 2014), suggesting that plant cell wall remodeling through pectin and cellulose modifications may be involved in Hartig net development. We previously reported that the symbiosis-induced ß-1,4-endoglucanase LbGH5-CBM1 from *L. bicolor*, acting on poplar cellulose, mannans and galactomannan, is involved in cell wall remodeling during formation of the Hartig net and this enzyme is required for successful symbiotic fungal colonization of poplar rootlets (Zhang *et al*., 2018). Also *L. bicolor* ß-glucanase GH131, active on polysaccharides with β-1,4, β-1,3, and mixed β-1,4/1,3 glucosidic linkages, is induced upon ectomycorrhiza formation (Anasontzis *et al*., 2019), further strengthening the concept of fungal PCWDEs as fundamental factors in the establishment of symbiotic plant-fungus interactions.

In the present study, we functionally characterized the symbiosis-upregulated *LbGH28A* from *L. bicolor* and confirmed its role in ectomycorrhiza formation by RNAi knock-down of the *LbGH28A* gene. A recombinant protein was produced and used for assaying the enzymatic activity of LbGH28A. These studies confirmed that LbGH28A is an endopolygalacturonase of family GH28. Finally, we localized the enzyme in *L. bicolor* cell walls and ectomycorrhizal interface by indirect immunofluorescence confocal microscopy and immunogold cytolocalization.

As showed by the phylogenetic analysis, this endopolygalacturonase is closely related to GH28A polygalacturonases from saprotrophic fungi. This suggests similar substrate specificities for these enzymes, namely activity on pectins in plant cell walls and plant residues, as well as a common evolutionary origin in Agaricomycotina. Transcription of *LbGH28A* in the free-living mycelium was induced by pectin, suggesting that its activity is indeed related to pectin metabolism. This was further confirmed by studies on the recombinant LbGH28A protein. The enzyme displays typical polygalacturonase activity and as expected, cannot hydrolyzes rhamnogalacturonans and galactans, which do not contain any polygalacturonic acid. Moreover, LbGH28A showed an optimal activity at pH 4 to 5, values similar to those of the plant apoplastic space. In addition, this degradative activity is enough to support the growth of *L. bicolor* mycelium on pectin as a sole carbon source (Fig. S1 in Veneault-Fourrey *et al*., 2014). Pectin methylation is an important factor influencing the activity of polygalacturonases (Lionetti *et al*., 2017). It remains to determine whether the symbiosis-induced pectin methylesterases (CE8) of *L. bicolor* act synergistically with LbGH28A to modify host pectins at the symbiotic interface.

Indirect immunofluorescence confocal microscopy using anti-LbGH28 antibodies showed that LbGH28A is localized at the periphery of *L. bicolor* hyphae forming the mantle sheath and the Hartig net. Its co-localization with/or adjacent to chitin label supports an extracellular/cell wall localization for this enzyme. Finally, immunogold cytolocalization microscopy confirmed that LbGH28A accumulates at the surface of both fungal and plant cells walls, i.e. the symbiotic interface, at the hyphal front penetrating between the epidermal root cells, while the LbGH5-CBM1 was mainly detected in the fungal cell walls.

Although *L. bicolor* lacks GH6 and GH7 cellobiohydrolases (Martin *et al*., 2008), the endocellulase LbGH5-CBM1 (Zhang *et al*., 2018), ß-glucanase GH131 (Anasontzis *et al*., 2019), expansins (Veneault-Fourrey *et al*., 2014) and here characterized polygalacturonase LbGH28A are likely involved in the limited alteration/remodelling of the host cell walls, thereby facilitating root penetration and Hartig net differentiation. We speculate that cellulose-acting LPMO (AA9) and GH5-CBM1 are involved in the loosening of the cellulose microfibrils and LbGH28A in remodeling the middle lamella through pectin (homogalacturonan) hydrolysis. However, it remains to be assessed how the mechanical cell wall properties, such as loosening and softening, and the strength and extension of adhesion areas between adjacent epidermal cells are altered during hypae colonization. These plant cell wall modifications may involve the depolymerization of matrix glucans and pectins, the pectin solubilization and removal of neutral sugars from pectin side chains. Such changes are expected to weaken the host cell walls and increase cell separation, which in combination with the turgor pressure of the hyphal tip, bring about textural changes. Atomic force microscopy (Zhang *et al*., 2016) will be used to further characterize the nanostructure of plant cell wall polysaccharides during ectomycorrhiza development. We speculate that these subtle modifications in the cell wall structures do not trigger the host plant immune response. Even if a moderate plant immune response is activated by cell wall elicitors, mycorrhiza-induced small effectors, such as the *L. bicolor* MiSSP7 and MiSSP7.6, would be able to dampen the defense reactions (Martin *et al*., 2016).

## Acknowledgments

The research was funded by the French Research National Agency (ANR-14-CE06-0020, project FUNTUNE), the Laboratory of Excellence Advanced Research on the Biology of Tree and Forest Ecosystems (ARBRE; ANR-11-LABX 0002 01, project SYMWOOD), the Genomic Science Program, US Department of Energy, Office of Science, Biological and Environmental Research as part of the Plant-Microbe Interfaces Scientific Focus Area (http://pmi.ornl.gov), and by grants from Universidad Nacional de Quilmes, Consejo Nacional de Investigaciones Científicas y Técnicas, and Agencia Nacional de Promoción Científica y Tecnológica; and by grants from National Natural Science Foundation of China (31901279), Natural Science Foundation of Gansu Province (20JR5RA276), and Fundamental Research Funds for the Central Universities (lzujbky-2019-ct01). The immunogold cytochemical electron microscopy was performed on the PiCSL-FBI core facility (IBDM, AMU-Marseille), a member of the France-BioImaging national research infrastructure (ANR-10-INBS-04). We would like to thank Dr. Raffaella Balestrini (University of Torino, Italy) for a gift of WGA.

## Author contributions

F.M. and J-G.B. planned and designed the research. F.Z., A.L., N.B.; M.H.; M.K., and A.D. performed experiments. F.Z., G.E.A., A.P., C.V-F., A.K., M-N.R., B.H., J-G.B. and F.M. analyzed the data. F.M. and F.Z. wrote the manuscript with the help of the co-authors.

---

**Supporting Information Table S1.**
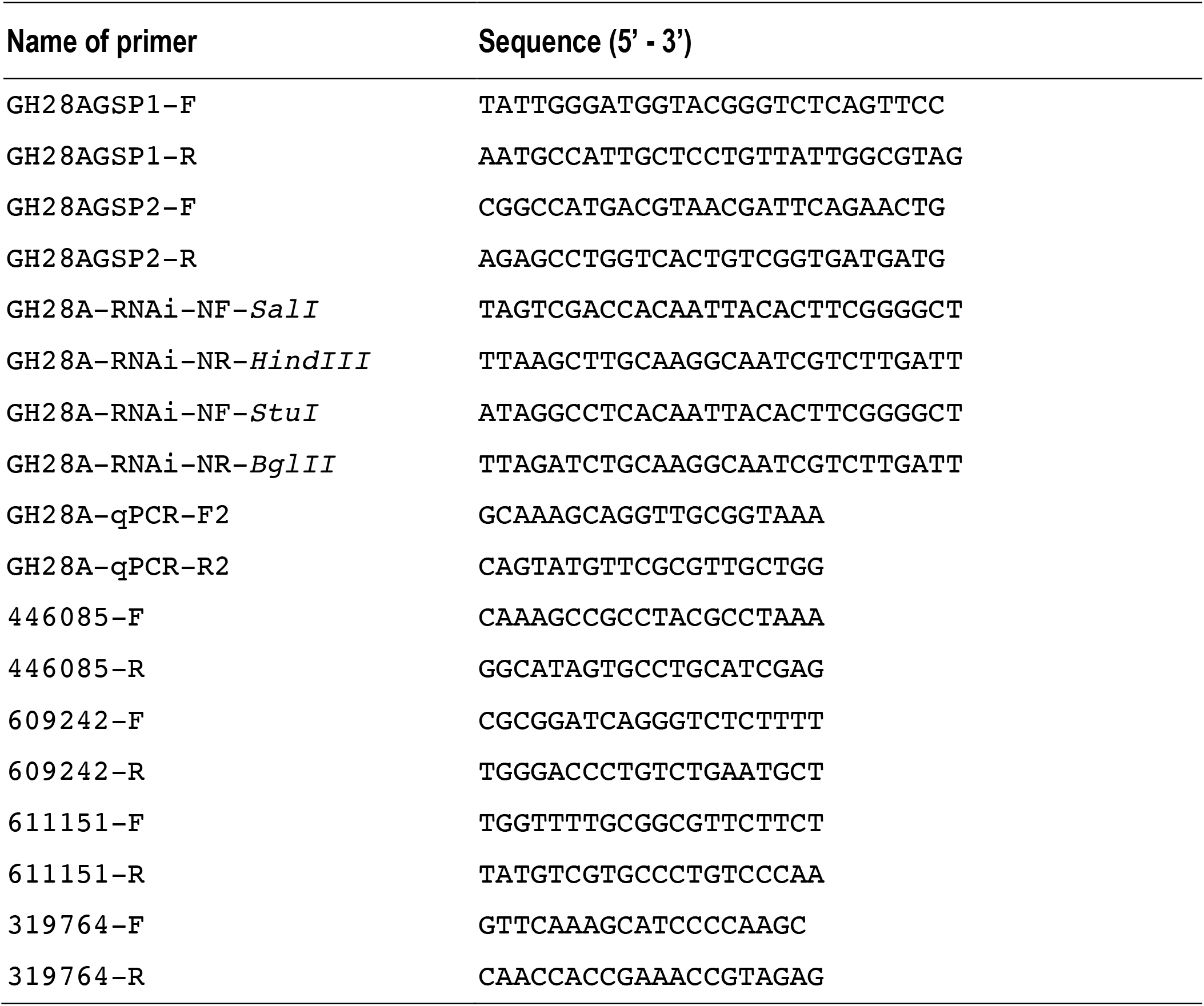
List of primer sequences used in this study.

## Supplementary Figures

**Figure S1.**
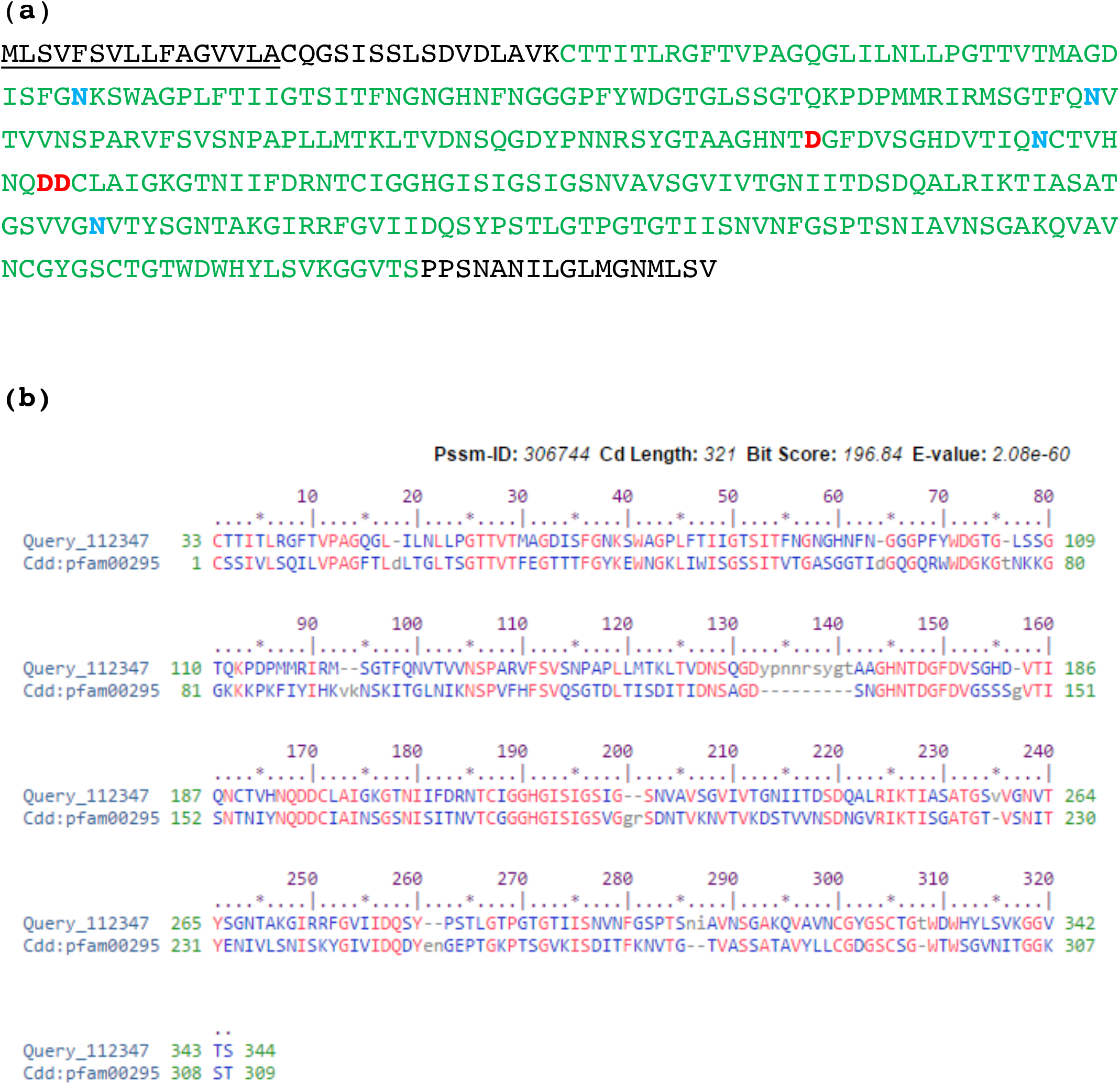
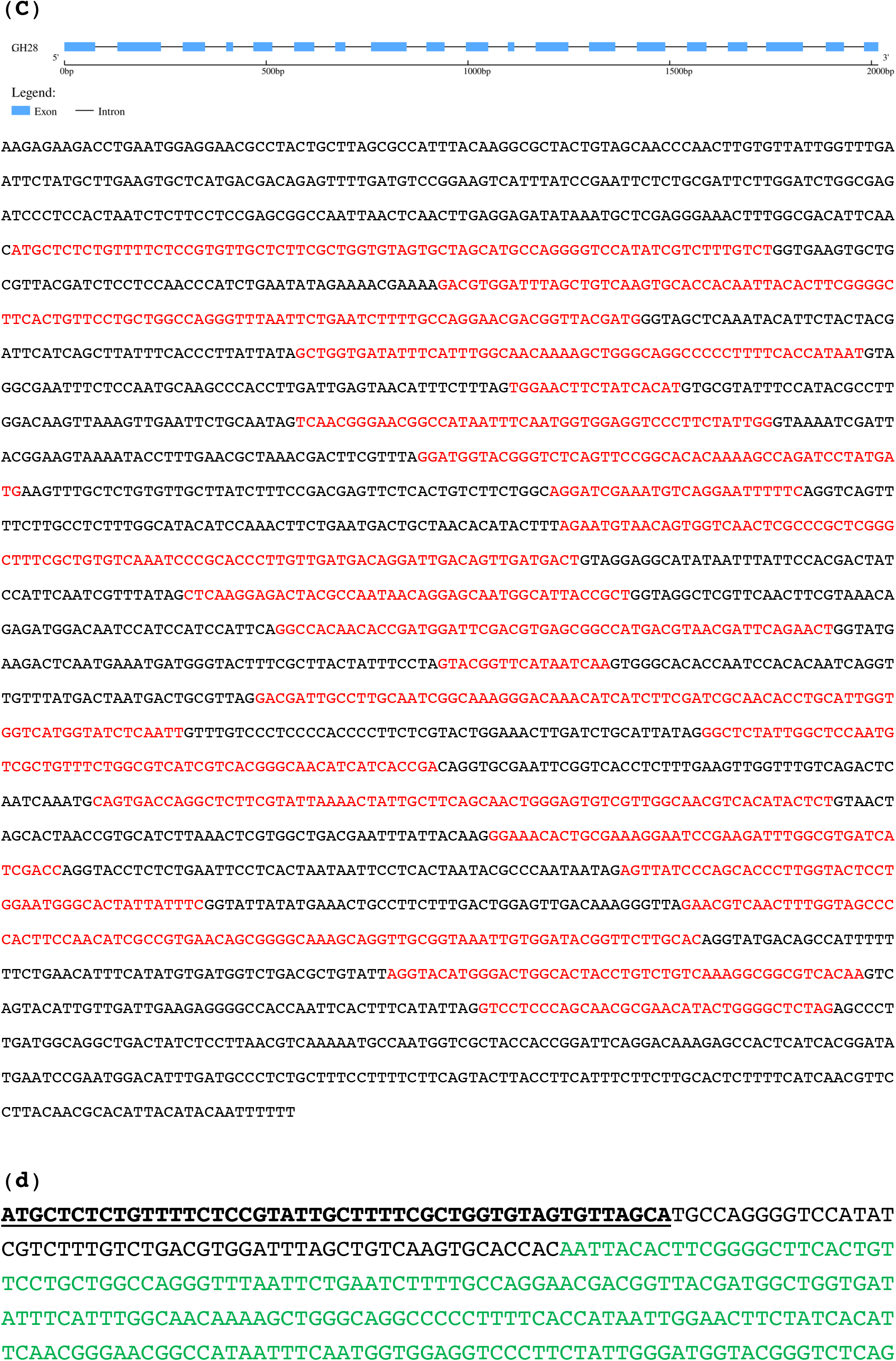
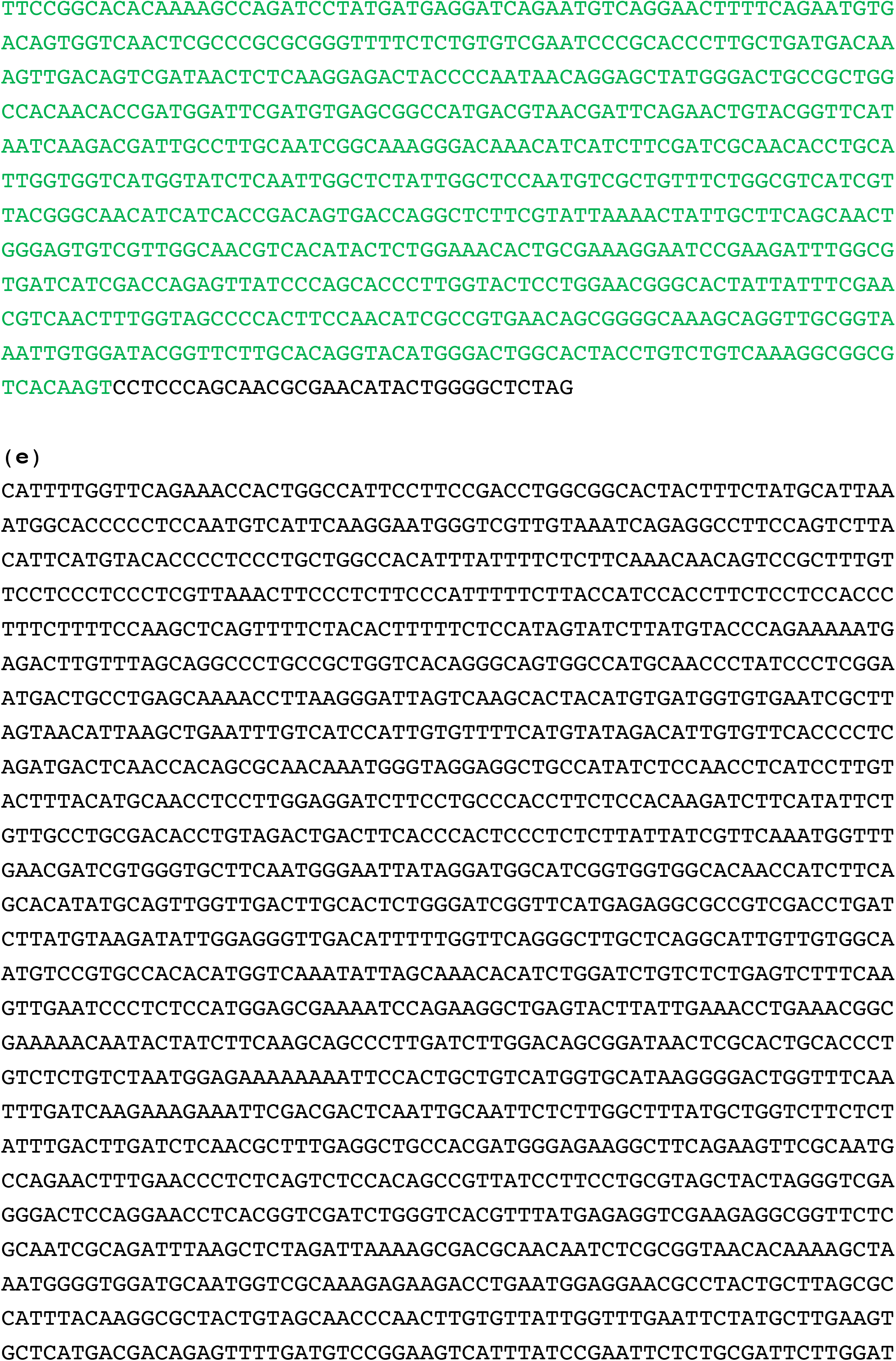
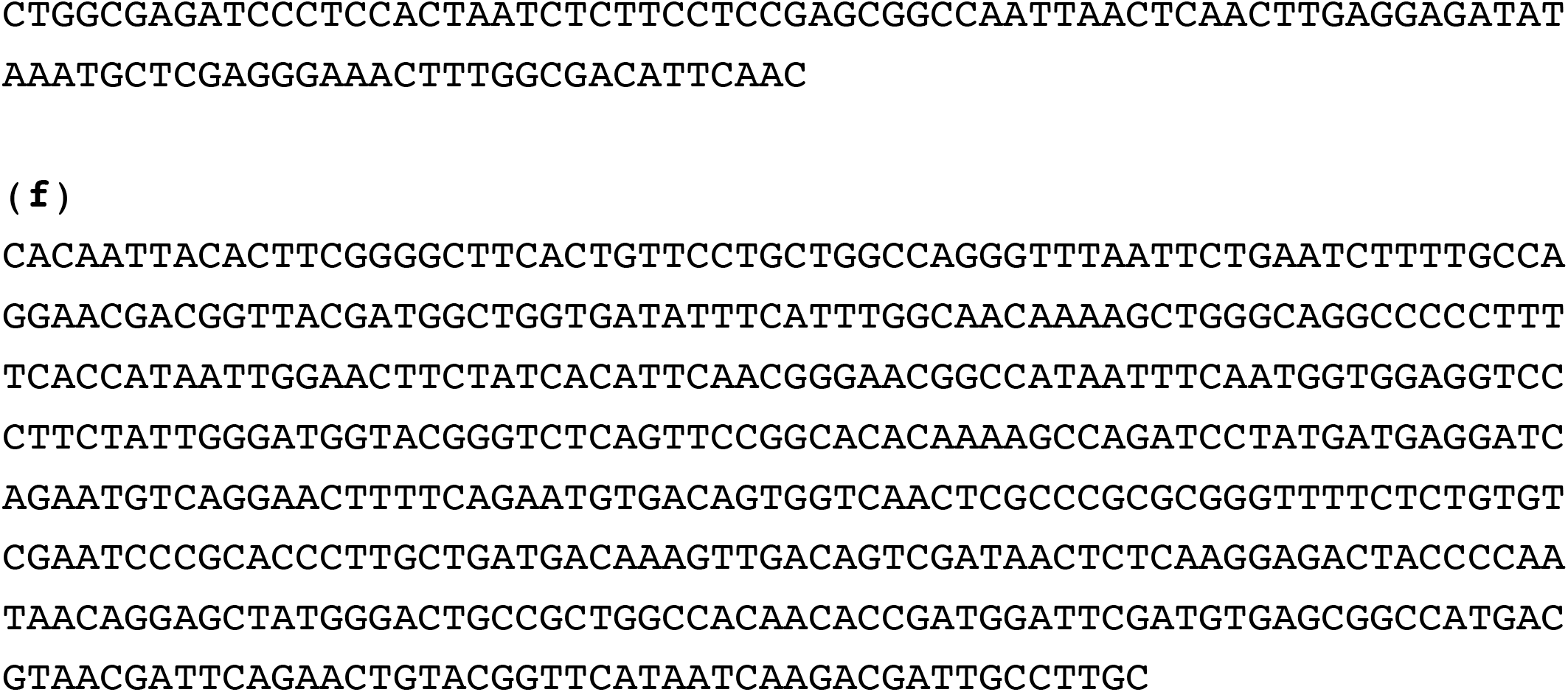
The *L. bicolor* GH28A polygalacturonase. (**a**) Amino acid sequence highlighting the native signal peptide (underlined), the putative N-glycosylation sites (in blue) and the conserved catalytic residues (in red); (**b**) PFAM domain (pfam00295) motif; (**c**) genomic sequence with 19 exons (in red) and 18 introns (in black); (**d**) coding (CDS) sequence; (**e**) nucleotide sequences of *LbGH28_A* promoter and (**f**) the nucleotide sequence of the RNAi target. Signal peptide is underlined and the catalytic domain is displayed in green.

**Figure S2.**
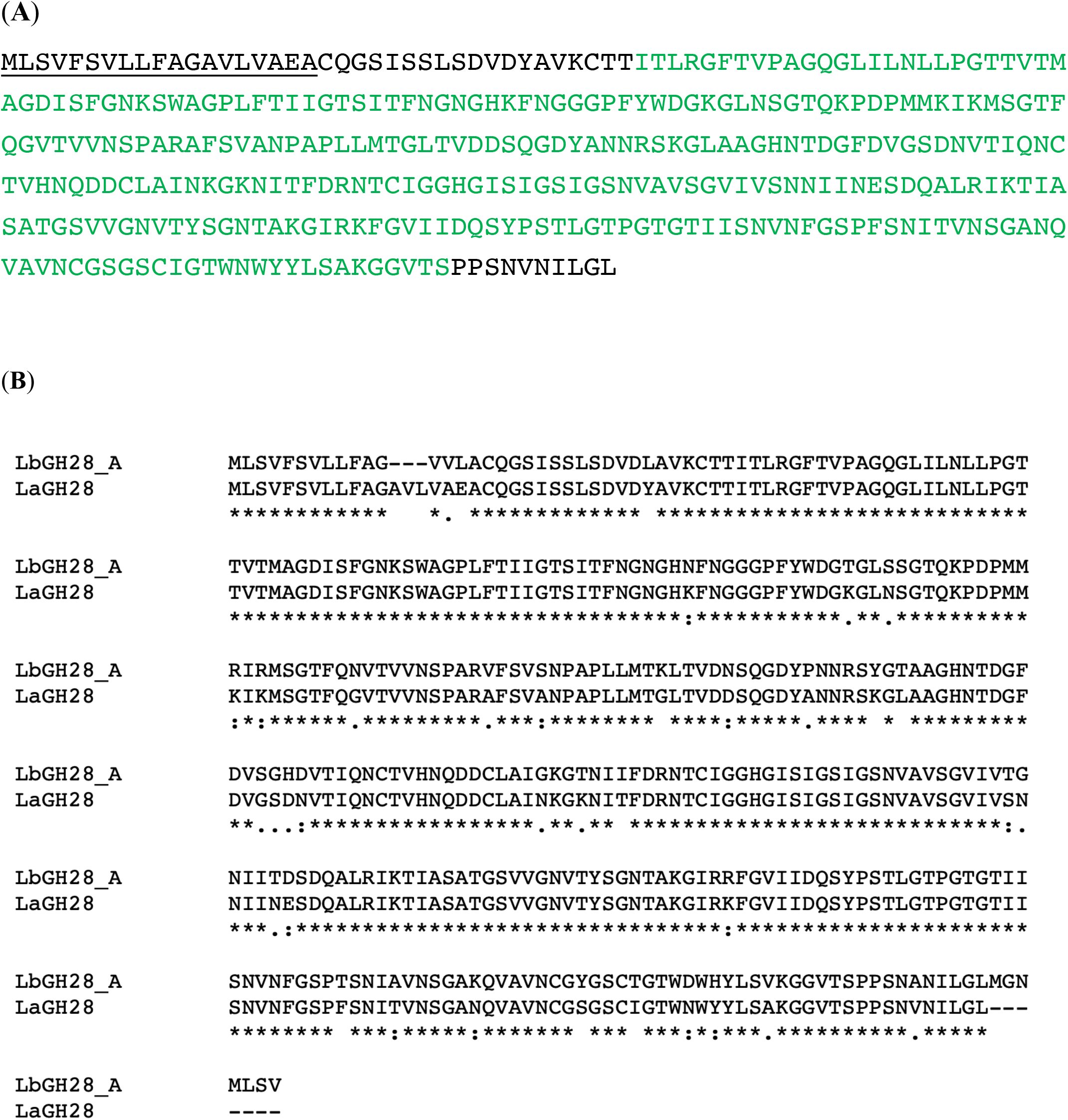
The *L. amethystina* GH28A polygalacturonase. (**A**) Amino acid sequence; (**B**) Alignment of *L. bicolor* and *L. amethystina* GH28A protein sequences.

## Notes

### Competing Interest Statement

The authors have declared no competing interest.

## References

Anasontzis GE, Lebrun M, Haon M, Champion C, Kohler A, Lenfant N, Martin F, O’Connell RJ, Riley R, Grigoriev IV et al. 2019. Broad-specificity GH131 β- glucanases are a hallmark of fungi and oomycetes that colonize plants. Environ Microbiol 21:2724–2739.

Balestrini R, Bonfante P. 2014. Cell wall remodeling in mycorrhizal symbiosis: a way towards biotrophism. Front Plant Sci 5:237.

Balestrini R, Kottke I. 2016. Molecular mycorrhizal symbiosis. John Wiley & Sons, Inc 47–61.

Balestrini R, Hahn MG, Faccio A, Mendgen K, Bonfante P. 1996. Differential localization of carbohydrate epitopes in plant cell walls in the presence and absence of arbuscular mycorrhizal fungi. Plant Physiol 111:203–213.

Brundrett MC. 2002. Coevolution of roots and mycorrhizas of land plants. New Phytologist 154:275–304.

Christiaens S, Van Buggenhout S, Houben K, Jamsazzadeh Kermani Z, Moelants KR, Ngouémazong ED, Van Loey A, Hendrickx ME. 2016. Process-structure-function relations of pectin in food. Crit Rev Food Sci Nutr. 56:1021–42.

Couturier M, Feliu J, Haon M, Navarro D, Lesage-Meessen L, Coutinho PM, Berrin JG. 2011. A thermostable GH45 endoglucanase from yeast: impact of its atypical multimodularity on activity. Microb Cell Fact 10:103.

Dietz S, von Bülow J, Beitz E, Nehls U. 2011. The aquaporin gene family of the ectomycorrhizal fungus Laccaria bicolor: lessons for symbiotic functions. New Phytologist 190:927–940.

Ellis SB, Brust PF, Koutz PJ, Waters AF, Harpold MM, Gingeras TR. 1985. Isolation of alcohol oxidase and two other methanol regulatable genes from the yeast Pichia pastoris. Mol. Cell. Biol 5:1111–1121.

Felten J, Kohler A, Morin E, Bhalerao RP, Palme K, Martin F, Ditengou FA, Legue V. 2009. The ectomycorrhizal fungus Laccaria bicolor stimulates lateral root formation in poplar and Arabidopsis through auxin transport and signaling. Plant Physiol 151:1991–2005.

Gay G, Normand L, Marmeisse R, Sotta B, Debaud JC. 1994. Auxin overproducer mutants of Hebeloma cylindrosporum Romagnesi have increased mycorrhizal activity. New Phytologist 128:645–657.

Gea L, Normand L, Vian B, Gay G. 1994. Structural aspects of ectomycorrhiza of Pinus pinaster (Ait.) Sol. formed by an IAA-overproducer mutant of Hebeloma cylindrosporum Romagnési. New Phytologist 128:659–670.

Kemppainen MJ, Pardo AG. 2010. pHg/pSILBAgamma vector system for efficient gene silencing in homobasidiomycetes: optimization of ihpRNA - triggering in the mycorrhizal fungus Laccaria bicolor. Microb Biotechnol 3: 178–200.

Kohler A, Kuo A, Nagy LG, Morin E, Barry KW, Buscot F, Canback B, Choi C, Cichocki N, Clum A et al. 2015. Convergent losses of decay mechanisms and rapid turnover of symbiosis genes in mycorrhizal mutualists. Nat Genet 47:410–415.

Koutz P, Davis GR, Stillman C, Barringer K, Cregg J, Thill G. 1989. Structural comparison of the Pichia pastoris alcohol oxidase genes. Yeast 5: 167–177.

Laurent P, Voiblet C, Tagu D, De Carvalho D, Nehls U, De Bellis R, Balestrini R, Bauw G, Bonfante P, Martin F. 1999. A novel class of ectomycorrhiza-regulated cell wall polypeptides in Pisolithus tinctorius. Mol Plant Microbe In 12:862–871.

Martin F, Aerts A, Ahren D, Brun A, Danchin EGJ, Duchaussoy F, Gibon J, Kohler A, Lindquist E, Pereda V et al. 2008. The genome of Laccaria bicolor provides insights into mycorrhizal symbiosis. Nature 452:88–92.

Martin F, Kohler A, Murat C, Balestrini R, Coutinho PM, Jaillon O, Montanini B, Morin E, Noel B, Percudani R et al. 2010. Perigord black truffle genome uncovers evolutionary origins and mechanisms of symbiosis. Nature 464:1033–1038.

Martin F, Kohler A, Murat C, Veneault-Fourrey C, Hibbett DS. 2016. Unearthing the roots of ectomycorrhizal symbioses. Nature Reviews Microbiology 14:760–773.

Massicotte HB, Peterson RL, Melville LH. 1989. Hartig Net structure of ectomycorrhizae synthesized between Laccaria bicolor (Tricholomataceae) and two hosts: Betula alleghaniensis (Betulaceae) and Pinus resinosa (Pinaceae). Am. J. Bot 76:1654–1667.

Miyauchi S, Kiss E, Kuo A, Drula E, Kohler A, Sánchez-García M, Morin E, Andreopoulos B, Barry KW, Bonito G, et al. 2020b. Large-scale genome sequencing of mycorrhizal fungi provides insights into the early evolution of symbiotic traits. Nature Communications 11:5125.

Pachlewski, R. 1967. “Mikotrofizm systemu korzeniowego,” in Zarys Fizjologii Sosny Zwyczajnej, eds S. Bialobok, and W. Zelawski, (Poznan: PWN Warszaw).

Peterson RL, Massicotte HB. 2004. Exploring structural definitions of mycorrhizas, with emphasis on nutrient-exchange interfaces. Canadian Journal of Botany 82:1074–1088.

Peter M, Kohler A, Ohm RA, Kuo A, Krützmann J, Morin E, Arend M, Barry KW, Binder M, Choi C. et al. 2016. Ectomycorrhizal ecology is imprinted in the genome of the dominant symbiotic fungus Cenococcum geophilum. Nat. Commun. 7, 12662.

Pitarch A, Sánchez M, Nombela C, Gil C. 2002. Sequential fractionation and two-dimensional gel analysis unravels the complexity of the dimorphic fungus Candida albicans cell wall proteome. Mol Cell Proteomics 1: 967–982.

Pfaffl, MW. 2001. A new mathematical model for relative quantification in real-time RT–PCR. Nucleic Acids Research 29:e45.

Read DJ, Leake JR, Perez-Moreno J. 2004. Mycorrhizal fungi as drivers of ecosystem processes in heathland and boreal forest biomes. Canadian Journal of Botany 82:1243–1263.

Smith SE, Read D. 2008. Mycorrhizal Symbiosis (Third Edition). London: Academic Press.

Tagu D, De Bellis R, Balestrini R, De Vries OMH, Piccoli G, Stocchi V, Bonfante P, Martin F. 2001. Immunolocalization of hydrophobin HYDPt-1 from the ectomycorrhizal basidiomycete Pisolithus tinctorius during colonization of Eucalyptus globulus roots. New Phytologist 149:127–135.

Tschopp JF, Brust PF, Cregg JM, Stillman CA, Gingeras TR. 1987. Expression of the lacZ gene from two methanol-regulated promoters in Pichia pastoris. Nucleic Acids Research 15: 3859–3876.

van der Heijden MG, Martin FM, Selosse MA, Sanders IR. 2015. Mycorrhizal ecology and evolution: the past, the present, and the future. New Phytol 205:1406–1423.

Veneault-Fourrey C, Commun C, Kohler A, Morin E, Balestrini R, Plett J, Danchin E, Coutinho P, Wiebenga A, de Vries RP et al. 2014. Genomic and transcriptomic analysis of Laccaria bicolor CAZome reveals insights into polysaccharides remodelling during symbiosis establishment. Fungal Genet Biol 72:168–181.

Zhang T, Zheng Y, Cosgrove DJ. 2016. Spatial organization of cellulose microfibrils and matrix polysaccharides in primary plant cell walls as imaged by multichannel atomic force microscopy. Plant J. 8:179–92.

Zhang F, Anasontzis GE, Labourel A, Champion C, Haon M, Kemppainen M, Commun C, Deveau A, Pardo A, Veneault-Fourrey C et al. 2018. The ectomycorrhizal basidiomycete Laccaria bicolor releases a secreted β-1,4 endoglucanase that plays a key role in symbiosis development. New Phytol. 220:1309–1321.

Zhang T, Tang H, Vavylonis D, Cosgrove DJ. 2019. Disentangling loosening from softening: insights into primary cell wall structure. Plant J. doi:10.1111/tpj.14519

